# *SNRK3.15* is a crucial component of the sulfur deprivation response in *Arabidopsis thaliana*

**DOI:** 10.1101/2025.04.29.651231

**Authors:** Anastasia Apodiakou, Elmien Heyneke, Saleh Alseekh, Pinnapat Pinsorn, Sabine Metzger, Stanislav Kopriva, Waltraud Schulze, Rainer Hoefgen, Sarah J. Whitcomb

**Affiliations:** Max Planck Institute of Molecular Plant Physiology, Am Mühlenberg 1, Potsdam-Golm, 14476 Germany; Institute for Plant Sciences, Cluster of Excellence on Plant Sciences (CEPLAS), University of Cologne, Zülpicher Str. 47b, Cologne, 50674 Germany; Department of Plant Systems Biology - University of Hohenheim, Garbenstraße 30, Stuttgart, 70599 Germany; Cereal Crops Research Unit, United States Department of Agriculture - Agricultural Research Service, Madison, WI, USA

**Keywords:** CIPK14/SNRK3.15 (AT5G01820), nutrient deficiency induced senescence (NuDIS), sulfate, chlorophyll, O-acetylserine (OAS)-cluster genes, proteomics

## Abstract

Sulfate deprivation (-S) results in numerous metabolic and phenotypic alterations in plants. Kinases are often key players in transducing nutrient status signals to molecular components involved in metabolic and developmental program regulation, but despite the physiological importance of sulfur, to date, no signaling kinases have been identified in sulfur-deficiency signaling response programs. Here we show that the serine/threonine protein kinase CIPK14/SNRK3.15 plays a regulatory role in the –S response in *Arabidopsis thaliana* seedlings. Multiple molecular and physiological responses to -S are attenuated in *snrk3.15* mutants, including both early adaptive responses and later emergency salvage processes including nutrient deficiency induced senescence. When grown in soil with sufficient sulfur supply, *snrk3.15* mutants showed no clear phenotypes, including no difference in seed sulfur content. Lastly, the proteome dataset generated from Col-0 and *snrk3.15.1* Arabidopsis seedlings under –S conditions for this project is the first of its kind and will be a valuable research resource. Proteomics data are available via ProteomeXchange with identifier PXD046612.

## INTRODUCTION

Plants cope with numerous abiotic and biotic stresses that negatively affect plant growth and development (Chaudhry & Sidhu, 2022). Exposure to abiotic stresses such as high or low temperature, drought or flood, soil salinity, and nutrient deficiency is detected by the plant, which activates responses on the molecular, cellular, and physiological levels (R. Li et al., 2009). One of these stress conditions is insufficient sulfur availability. Plants primarily rely on sulfate in the soil as their sulfur source for biosynthesis of a broad range of metabolites that play essential roles in plant growth and metabolism (V. J. Nikiforova et al., 2004). Cysteine (Cys) is the first reduced organic sulfur compound, which integrates the sulfur, nitrogen, and carbon assimilation pathways (Jobe et al., 2019). The sulfur-containing amino acids Cys and methionine (Met) are not only incorporated into proteins during translation but are also precursors of compounds such as glutathione (GSH), plant defensins such as glucosinolates (GSL), and cofactors such as acetyl-CoA (Hoefgen & Hesse, 2015; Takahashi et al., 2011). GSH plays an important role in redox regulation and facilitates the detoxification of reactive oxygen species and heavy metals (Shanmugam et al., 2015). Acetyl-CoA is oxidized in the TCA cycle, resulting in the production of the reducing equivalents NADH and FADH2, which transfer electrons to the mitochondrial respiratory chain (Oliver et al., 2009). It is also a building block in the synthesis of fatty acids, amino acids, and glucosinolates (Field et al., 2004). Acetyl-CoA is needed to synthesize the direct precursor of Cys, O-acetylserine (OAS), from serine (Hoefgen & Hesse, 2015). Met is the precursor of S-adenosylmethionine (SAM). SAM is vital for plants since it is a central methyl donor for numerous methylation reactions, including those in chlorophyll biosynthesis and on chromatin (Loenen, 2006). Additionally, proteins containing iron-sulfur (Fe-S) clusters are redox-active cofactors and enzymes essential for many core metabolic processes, including photosynthesis, cellular respiration, and DNA metabolism (Fonseca et al., 2021; Zuchi et al., 2015). Hence, sulfur is a crucial nutrient in primary and secondary metabolism and is ultimately important for plant growth and performance (Hoefgen & Hesse, 2007).

Sulfate deficiency leads initially to the reduction of sulfur containing metabolites such as GSH, GSL, and SAM, while ultimately resulting in Cys, and Met reduction (Aarabi et al., 2020; Forieri et al., 2017; V. J. Nikiforova et al., 2005, 2006). Visible responses to sulfur starvation include growth retardation, accumulation of anthocyanins, and chlorosis (Forieri et al., 2017; J. Zhang et al., 2011; V. J. Nikiforova et al., 2004). Anthocyanins are a class of flavonoids with antioxidant properties that may help to mitigate the effects of stress-induced elevated ROS levels, especially in a context where GSH levels cannot be maintained due to insufficient S-supply. SAM decreases under -S, likely contributing to chlorosis since SAM is the methyl donor for methylation reactions in the last step of chlorophyll biosynthesis (Hawkesford et al., 2011; V. J. Nikiforova et al., 2005, 2006). Shoot growth is typically reduced to a much greater extent than root growth, resulting in an increased root:shoot ratio under –S. Root system architecture is also altered under -S, however S-starvation responses differ among studies. Kutz and colleagues reported that – S stimulates lateral root initiation via NITRILASE 3 (NIT3) (Kutz et al., 2002). Similar to Kutz 2002, Nikiforova and colleagues reported that –S conditions result in extensive root elongation and branching, perhaps as an adaptation to search for sulfur sufficient areas (V. Nikiforova et al., 2003). It was further reported that plants under –S develop increased root branching with denser lateral roots closer to the root tip (López-Bucio, Cruz-Ramírez & Herrera-Estrella, 2003). In contrast to this, two studies showed inhibition of lateral root initiation and stimulation of primary root growth under –S (Joshi et al., 2019; Hubberten et al., 2012), and other studies showed that sulfur deprivation of seedlings resulted in decreased lateral root density (Dong et al., 2019). It has even been reported that sulfur starvation of seedlings did not lead to altered root architecture at all (Gruber et al., 2013). The lack of consistency in how root system morphology responds to S-deficiency has been speculated to be due to plant age during the −S treatment (Gruber et al., 2013). Moreover, we assume that physicochemical differences in growing parameters and composition of the growth media additionally provoked these differing observations (Hannah et al., 2010).

When plants are shifted to –S, sulfide availability decreases, leading to the accumulation of OAS and induction of the OAS-cluster genes: *SULFUR DEFICIENCY INDUCED* genes 1 and 2 (*SDI1* and *SDI2*), *MORE SULFUR ACCUMULATION1* (*MSA1*), *GAMMA-GLUTAMYL CYCLOTRANSFERASE 2;1* (*GGCT2;1*), *APS REDUCTASE 3* (*APR3*), and *RESPONSE TO LOW SULFUR 1* (*LSU1*) (Hubberten et al., 2012). SDI1 and SDI2 inhibit accumulation of GSLs, sulfur-rich seed storage proteins, and sulfolipids (Aarabi et al., 2023, 2021, 2016). MSA1 is a regulator of SAM biosynthesis and DNA methylation (Huang et al., 2016), and when plants are grown under –S, GGCT2;1 has been shown to contribute to GSH degradation via the γ-glutamyl cycle, most likely to mobilize Cys (Joshi et al., 2019). APR3 is localized in the chloroplasts, where it participates in the reduction of adenosine 5’-phosphosulfate (APS) to sulfite (Koprivova et al., 2008). Lastly, LSU1 has been hypothesized to be involved in autophagy under –S conditions (Sirko et al., 2015) and together with other members of the *LSU* gene family act as modulators of the –S response (Piotrowska et al., 2024).

It is well established that nutrient signaling depends upon conserved kinases, such as TARGET OF RAPAMYCIN (TOR), SUCROSE NON-FERMENTING-RELATED KINASE 1 (SNRK1) (Robaglia et al., 2012), and SNRK3 family kinases (Yasuda et al., 2017). However, despite the physiological importance of sulfur, mechanisms of sulfur signaling remain elusive. Among the 26 SNRK3 kinases in Arabidopsis, *SNRK3.15* (*AT5G01820*), also known as *CBL-INTERACTING PROTEIN KINASE 14* (*CIPK14*) and *SALT RESPONSIVE 1* (*SR1*), is among the most strongly induced under sulfur deprivation conditions (Yuan et al., 2020; Jonathan W. Mueller & Naeem Shafqat, 2015; Iyer-Pascuzzi et al., 2011; Maruyama-Nakashita et al., 2003). And although involvement in S-nutrient signaling has yet to be shown (Heyneke et al., 2015), DAP-seq and RNA-seq (Dietzen et al., 2020; O’Malley et al., 2016) data suggest that *SNRK3.15* expression is directly regulated by SULFUR LIMITATION 1 (SLIM1), a key transcription factor (TF) regulating S-starvation responsive genes under –S conditions. SNRK3 kinases are activated via interaction with Ca^2+^ sensing calcineurin B-like proteins (CBL) (Weinl & Kudla, 2009). The CBL-SNRK3 signaling pathway is known to crosstalk with the ABA signaling pathway (Yu et al., 2014), and ABA is known to decrease under –S conditions (Cao et al., 2014). Furthermore, CBL-SNRK3 complexes have been shown to directly regulate the transport of potassium and nitrate across the plasma membrane via phosphorylation of the transporters NRT1.1 (nitrate low affinity transporter) and AKT1 (K^+^ transporter) (Ho et al., 2009; Xu et al., 2006). Whether sulfate transporter activity is regulated *in planta* by phosphorylation is unknown, but it has been shown in a heterologous system that a putative phosphorylation site in the STAS domain of SULTR1;2 is necessary for sulfate transport (Rouached et al., 2009). A *SNRK3.15* mutant (*snrk3.15.1*, SALK_009699) showed stronger growth than Col-0 under low phosphate conditions, indicating the involvement of SNRK3.15 in growth regulation under phosphate deprivation (Linn et al., 2017). However, to our knowledge, no previous reports have shown a SNRK3 kinase to be functionally involved in sulfur deficiency responses or growth regulation.

## MATERIALS AND METHODS

### Plant material

*Arabidopsis thaliana* Columbia-0 (Col-0), and the T-DNA mutant lines SALK_009699 *(snrk3.15.1*) and SALK_147899 *(snrk3.15.2*) were obtained from the Nottingham Arabidopsis Stock Centre (University of Nottingham). The T-DNA insertion position of both mutants has been reported previously (Lin et al., 2014; Qin et al., 2008). In both SALK_009699 and SALK_147899, the T-DNA is inserted in the sole exon of the SNRK3.15 gene (AT5G01820) (Figure S1a). Primers for genotyping were designed using the SALK T-DNA tool (http://signal.salk.edu/tdnaprimers.2.html) with Lb13 as T-DNA-specific primer (Supplemental Table 1).

### Plant cultivation

Agar plates: *Arabidopsis thaliana* seeds were sterilized using chlorine gas, prepared by combining 100 ml of NaClO and 5 ml of HCl. The seeds were then sown on 750 μM MgSO_4_ FN sterile agar medium, as described in Supplemental Table 2, and were stratified at 4 °C for two nights before the plates were placed vertically in controlled environment chambers (CLF Plant Climatic). The seeds germinated under a 16 h photoperiod with an irradiance of 100 µmol photons m^−2^ s^−1^. The temperature inside the chambers was maintained at 21 °C during the day and 19 °C at night. After 5 days [7 days after sowing (DAS)] the seedlings were carefully transferred with tweezers to fresh FN or –S plates, and they were returned into the growth chamber, vertically. After 11 days (18 DAS), photos of the seedlings were taken.

Liquid shaking cultures: *Arabidopsis thaliana* seeds (3 mg) were sterilized with the chlorine gas method and were then put in sterile 250 ml glass flasks containing 30 ml of 300 μΜ MgSO_4_ FN liquid media, as described in Supplemental Table 2. After 2 nights of stratification, the flasks were transferred to a constant light growth chamber with mild shaking at 75 rpm. After 7 days of growth under light (9 DAS), the seedling ball was rinsed with sterile ddH2O to remove excess MgSO_4_, and the seedling ball was transferred to either fresh FN or –S media [0 days after transfer (DAT)]. The plants were returned to light and harvested at 1 DAT, 3 DAT, and 7 DAT. Immediately prior to snap-freezing in liquid nitrogen, the seedling ball was rinsed with sterile ddH2O and well-dried. The frozen tissue was homogenized using a Retschmill MM400.

Greenhouse: Seedling that were not transferred to fresh FN or –S agar plates, as described in the agar plates subsection above, were instead transplanted into soil-filled round pots with a diameter of 6 cm. These pots were placed in a growth chamber with controlled conditions: photoperiod of 16 hours with an irradiance of 120 µmol photons m^−2^ s^−1^, temperature of 20 °C during the day and 16°C at night, and a humidity level of 60-75%. To minimize the potential effects of uneven lighting, the positions of the pots were regularly rearranged within the growth chamber.

### Determination of leaf area, root architecture, plant biomass

For leaf area and root architecture measurements on agar plate-grown seedlings, photos were taken with a Keyence VHX6000 microscope. To facilitate subsequent image processing, a scale was included in the photos, and image analysis software, ImageJ (National Institute of Health, USA), was used for further analysis. For the rosette area measurements of growth chamber-grown plants in soil, photographs were taken from above once a week using a professional camera. The distance between the pots and the camera was kept consistent throughout each experiment. The total rosette area was determined from the photos using the image analytic software, ImageJ (National Institute of Health, USA). For biomass measurements, seedling balls from shaking cultures were carefully dried using tissue paper and the fresh weight was determined using an electronic semi-microbalance (VWR).

### Seed weight

Plants grown in the greenhouse under photoperiod of 16h (see plant cultivation section, above) that were used for rosette area measurements, were allowed to grow in the growth chamber until they reached the stage of seed production. The plants were individually bagged, seeds allowed to mature for 1 month at 50% humidity and 20 °C day-/16 °C night-temperature, and then seeds were dried for 2 weeks in a drying room at 15 °C with 15% humidity. The dry seed weight was determined as follows: 50 seeds were collected at the end of the maturation period. After 2 additional weeks of drying, the dry weight of 50 seed pools was determined and the mean single seed mass was calculated.

### Chlorophyll and anthocyanin quantification

20 mg of homogenized tissue was suspended in 300 μl of ice-cold 95% (v/v) ethanol by briefly vortexing and kept on ice until all the samples were processed. To remove debris, the samples were centrifuged at 14000 x g for 5 min. The supernatant was carefully transferred to a new tube, which was then placed on ice and covered with foil to prevent chlorophyll degradation. For absorbance determination, 100 μl of the supernatant was brought to 200 μl with 95% (v/v) ethanol and put in a flat bottom 96-well plate. Absorbance measurements were taken at wavelengths of 664.1 nm, 648.6 nm, 470 nm, and 750 nm (blank). Chl a, and Chl b concentrations were calculated using the following formulas (Lichtenthaler & Buschmann, 2001), and calculated concentrations were normalized to the fresh weight (FW) of the extracted sample.

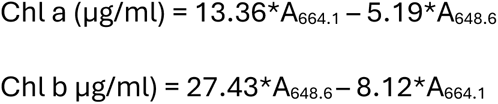

For anthocyanin content measurements, 90 μl of the chlorophyll-containing supernatant was mixed with 10 μl of 1M HCl, and the absorbance was measured at 520 nm and 750 nm (blank) (Lee et al., 2008). Absolute absorbance was normalized to the FW of the extracted sample.

### Glucosinolate and sulfate determination

25 mg of homogenized tissue was extracted as previously described with few modifications (Lisec et al., 2006) . Ice-cold CHCl_3_/CH_3_OH (3:7, v/v) supplemented with Isovitexin and ^13^C6-sorbitol for internal standards was added to the frozen homogenized tissue. The samples were placed at -20 °C for 2 h and vortexed briefly every 30 min. Ice-cold water was added, and samples were vortexed until the two phases were dispersed. Samples were centrifuged at 14000 x g for 10 min at 4 °C. The upper, polar phase was transferred to a new tube and desired volumes were further aliquoted for LC-MS (200 μl) and ion chromatography. The non-polar phase was also transferred to a new tube. The polar phase aliquots, non-polar phase, and insoluble material were evaporated using a centrifugal vacuum dryer at 30 °C for 5 h. For GSL determination, LC-MS was performed essentially as previously described (Perez de Souza et al., 2021, 2019). For anion content determination, the evaporated polar phase was dissolved in high purity H_2_O and analyzed by a Dionex ICS-3000 system using a 17-minute KOH gradient (6 mM - 55 mM, 0.25 ml/min flowrate). Sulfate concentration in the samples was calculated based on the known concentration of the following standards: (NH_4_)_2_SO_4_, MgSO_4_. The standards had a concentration range between 2 μM – 200 μM.

### Thiol determination

Extraction was performed as previously described (Guo, 2018; Watanabe et al., 2008). Briefly, 25 mg of homogenized tissue was mixed with 0.1 M HCl, and samples underwent a second round of homogenization using a Retschmill to achieve optimal cell lysis. For the reduction step with 1.58 M *N*-ethylmorpholine, the extract was supplemented with 25 μN-acetyl-Cys, as the internal standard. The reaction was allowed to react with 25 mM phosphine for 20 min at 37 °C. For the labeling step, the reduced sample reacts with 30 mM monobromobimane (mBrB) for 20 min at 37 °C in the dark. The labeling reaction was terminated by the addition of acetic acid, and the resulting solution was then subjected to HPLC analysis. HPLC was performed as previously described (Watanabe et al., 2008).

### OAS determination

O-acetylserine (OAS) was extracted as previously described (Hubberten et al., 2012). Briefly, 80% ethanol (EtOH) containing 2.5 mM HEPES at pH 6.2 was added to 35 mg of homogenized tissue sample. The samples were centrifuged at 14000 rpm for 10 min at 4°C. The resulting supernatant was transferred to a new tube, while the pellet was re-extracted using 50% EtOH/2.5 mM HEPES at pH 6.2, followed by shaking for 20 min at 4 °C. After centrifugation, as described above, the supernatant was collected and combined with the initial supernatant. The pellet was then extracted a final time with 80% EtOH, and the supernatant was added to the combined supernatants. The supernatants were shaken for 20 min at 4 °C, followed by centrifugation and the supernatant was subjected to HPLC analysis as described (Hubberten et al., 2012).

### Element analysis

Element concentrations were determined by inductively coupled plasma mass spectrometry (ICP-MS) according to the method described in (Almario et al., 2017).

Samples were dried before digestion. The dried material was ground to fine powder and 5 mg were digested in 15 ml Falcon tubes using 500 µL of HNO_3_ (67%) overnight at room temperature. The next day, loosely closed samples were placed in 95°C water bath until the liquid was completely clear (30 min). After cooling to room temperature for 10-15 min samples were put on ice and 4.5 ml deionized water was carefully added to the tubes and the tubes were weighed. The final solutions were centrifuged at 4°C at 4000 rpm for 30 min and supernatants were transferred to new tubes. The elemental concentration was determined using an Agilent 7700 ICP-MS (Agilent Technologies) following the manufacturer’s instructions. The dilution factor was calculated as follows: DF= (final weight – empty falcon weight) / sample DW.

### RNA extraction, cDNA synthesis, and qRT-PCR

Total RNA was isolated from the seedling ball using Sigma Spectrum Plant total RNA KIT (1003037777) according to the manufacturer’s instructions with on-column DNase treatment. The RNA concentration and purity were verified using an Agilent Bioanalyzer 2100. First-strand cDNA was synthesized from 1000 ng of total RNA using the PrimeScript II 1st Strand cDNA Synthesis Kit (TAKARA) according to the manufacturer’s instructions. The reactions were diluted (1:9), and 0.5 μl were used to perform RT-PCR reactions with SYBR Green PCR Master Mix (Applied Biosystems). The reaction set up and thermal program profile are provided in Supplemental Table 3. Data were analyzed using SDS 2.0 software (Applied Biosystems). C_t_ values for all genes were normalized to the geometric mean of the C_t_ values of *TIP41* and *PP2A*. NormFinder (Andersen et al., 2004) was used to select *TIP41* and *PP2A* as normalization genes because they show suitably stable expression in our given sample set and experimental design. Oligonucleotide primers for qPCR were designed using the NCBI primer designing tool (https://www.ncbi.nlm.nih.gov/tools/primer-blast/) and are provided in Supplemental Table 1.

### Proteomic profiling and data analysis

Proteomic profiling was performed on Col-0 and *snrk3.15.1* shaking culture samples from 3 DAT to FN and 3 DAT to -S, three replicates per sample group (genotype x condition). Tissue from three flasks was pooled for each replicate. Soluble proteins and microsomal membranes were isolated in the presence of phosphatase and protease inhibitors (50 mM NaF, 1 mM NaVO_4_, 4 μM leupeptin, 1 mM benzamidine, 0.03 μM microcystin). Extracted protein was enriched by precipitation and resuspended in 6 M urea, 2 M thiourea, pH 8.0 with TRIS-HCl. Protein was then pre-digested for 3 h with endoproteinase Lys-C (0.5 μg μl^-1^; Wako chemical, Neuss, Germany) at room temperature. Samples were diluted 1:4 with 10 mM TRIS-HCl (pH 8.0) and digested with 4 μl sequencing-grade modified trypsin (0.5 μg μl^-1^; Promega, Fichtburg, WI) at 37 °C, overnight.

Trifluoracetic acid was used to stop the digestion by acidifying the protein solution to < pH 3.0. Digested peptides were desalted over Stop And Go Extraction Tips (C18 Empore Disks) (Ishihama et al., 2006) and dissolved in 80% acetonitrile and 0.1% trifluoracetic acid. LC-MS/MS analysis was performed using a nano-flow HPLC system (Thermo Fisher Scientific, Dreieich, Germany) with a C18 analytical column (75 μm ID, 15 cm length) coupled to an LTQ-Orbitrap hybrid mass spectrometer (Thermo Scientific, Dreieich, Germany). Chromatography was performed with solution A (0.5% acetic acid) and solution B (0.5% acetic acid, 80% acetonitrile) using the following gradient: 5% solution B to 30% solution B over 71 min, then to 60% solution B over 14 min, and then to 90% solution B over 10 min). The eluted peptides were sprayed directly into the LTQ-Orbitrap mass spectrometer. The fragmentation spectra (of the multiple-charged peptides that were assimilated) were used to identify the peptides. Five MS spectra were acquired for each full scan spectrum, 60,000 full-width at half-maximum resolution. The mass spectrometry proteomics data have been deposited in the ProteomeXchange Consortium via the PRIDE (Deutsch et al., 2017) partner repository with the dataset identifier PXD046612.

Protein identification and ion intensity quantitation was carried out by MaxQuant version 1.5.3.8 (Cox & Mann, 2008). Spectra were matched against the Arabidopsis proteome (TAIR10, 35386 entries) using Andromeda (Cox et al., 2011). Thereby, carbamidomethylation of cysteine was set as a fixed modification; oxidation of methionine was set as variable modifications. Mass tolerance for the database search was set to 20 ppm on full scans and 0.5 Da for fragment ions. Multiplicity was set to 1. For label-free quantitation, retention time matching between runs was chosen within a time window of two minutes. Peptide false discovery rate (FDR) and protein FDR were set to 0.01, while site FDR was set to 0.05. Hits to contaminants (e.g. keratins) and reverse hits identified by MaxQuant were excluded from further analysis. Label-free quantitation based on LFQ values were used for quantitative analysis.

Peptide ion intensity values were normalized to the sum ion intensity value in the respective sample (Supplemental Table 4). Missing value imputation was performed on normalized intensity data with missForest v 1.5 R package (Stekhoven & Bühlmann, 2012) on microsomal and soluble fraction samples separately. Missing values for proteins detected in 5 or fewer samples were not imputed. Among proteins detected in at least 6 samples, 13% and 10% of values were imputed in microsomal and soluble samples respectively (Supplemental Table 4). Normalized intensity values were log2 transformed prior to differential abundance testing. Differential abundance testing was performed separately on data from microsomal and soluble fraction samples. Welch test p-values were adjusted by Benjamini-Hochberg method, and proteins with adjusted p-values < 0.1 were considered differentially abundant proteins (DAPs) (Supplemental Table 5).

Proteins were annotated with Gene Ontology terms using BiomaRt v2.56.1 R package (Durinck et al., 2009, 2005) to access the EnsemblPlants database (https://plants.ensembl.org) and with KEGG pathways and Enzyme Commission numbers using KEGGREST v1.40.0 R package (Tenenbaum & Maintainer, 2023) to access the Kyoto Encyclopedia of Genes and Genomes database (https://www.kegg.jp). Databases were accessed in November 2023. Gene Ontology annotations for S-responsive DAPs in Col-0 are provided in Supplemental Table 6. Over representation analysis was performed with the enricher() function from clusterProfiler v4.0 R package (Wu et al., 2021), with minGSSize = 5 and maxGSSize = 500 (Supplemental Table 7).

## Supplemental Data

Supplemental Figure 1: Supplemental characterization of *snrk3.15* lines, *SNRK3.15* expression, and comparative growth in soil

Supplemental Figure 2: Phenotypes of dry seed from Col-0 and *snrk3.15* plants grown on soil

Supplemental Figure 3: Response of chlorophyll and chlorophyll degradation genes to -S in Col-0 and *snrk3.15* seedlings

Supplemental Figure 4: Levels of proteins positively associated with chlorophyll content

Supplemental Figure 5: Levels of proteins negatively associated with chlorophyll content

Supplemental Figure 6: Exploratory analysis of global proteome

Supplemental Figure 7: Levels of -S responsive, *snrk3.15*-specific DAPs

Supplemental Figure 8: Relationship between sulfate, OAS, and OAS-cluster genes

Supplemental Table 1: Primers used for genotyping and gene expression

Supplemental Table 2: Media composition

Supplemental Table 3: qRT-PCR parameters

Supplemental Table 4: Proteome data

Supplemental Table 5: Welch-test results for condition-responsive and for genotype-responsive DAPs

Supplemental Table 6: Gene annotations for S-responsive DAPs in Col-0

Supplemental Table 7: Over representation analysis results for S-responsive DAPs in Col-0 and/or *snrk3.15.1*

## RESULTS

### *snrk3.15* seedlings are more tolerant of sulfur deficiency conditions than Col-0

As previously shown, SNRK3.15 is a kinase transcriptionally induced by -S conditions (Figure 1a) (Dietzen et al., 2020; Iyer-Pascuzzi et al., 2011; Maruyama-Nakashita et al., 2003; V. Nikiforova et al., 2003). To assess whether SNRK3.15 has a role in S-deficiency responses, growth of two independent *snrk3.15* knock-down mutants (Supplemental Figure 1a, b) was compared to the parental accession, Col-0. When grown on –S agar plates, the *snrk3.15* mutants displayed 1.7-fold greater total leaf area than Col- 0, while no differences in leaf area were found between Col-0 and *snrk3.15* on full nutrition (FN) plates (Figure 1b,c). When grown in soil, representing a FN state, there was no significant difference in the rosette leaf area between *snrk3.15* mutants and Col-0 (Supplemental Figure 1c,d). Under -S, differences in root architecture between Col-0 and *snrk3.15* mutants were observed. Col-0 responded to -S by increasing lateral root density while this did not occur in *snrk3.15* mutants (Figure 1d). On FN plates, Col-0 and *snrk3.15* displayed similar lateral root density. *NITRILASE 3* (*NIT3*) has been proposed to drive root proliferation under −S conditions and is expressed in lateral root primordia (Kutz et al., 2002). Nitrilases are membrane bound proteins (Bartling et al., 1994), therefore we checked NIT3 protein content in the microsomal protein fraction by LC-MS/MS and found that NIT3 protein accumulated in response to –S in both Col-0 and *snrk3.15.1*, but under -S NIT3 content in *snrk3.15.1* was 41% less than in Col-0 (Figure 1e).

**Figure 1:**
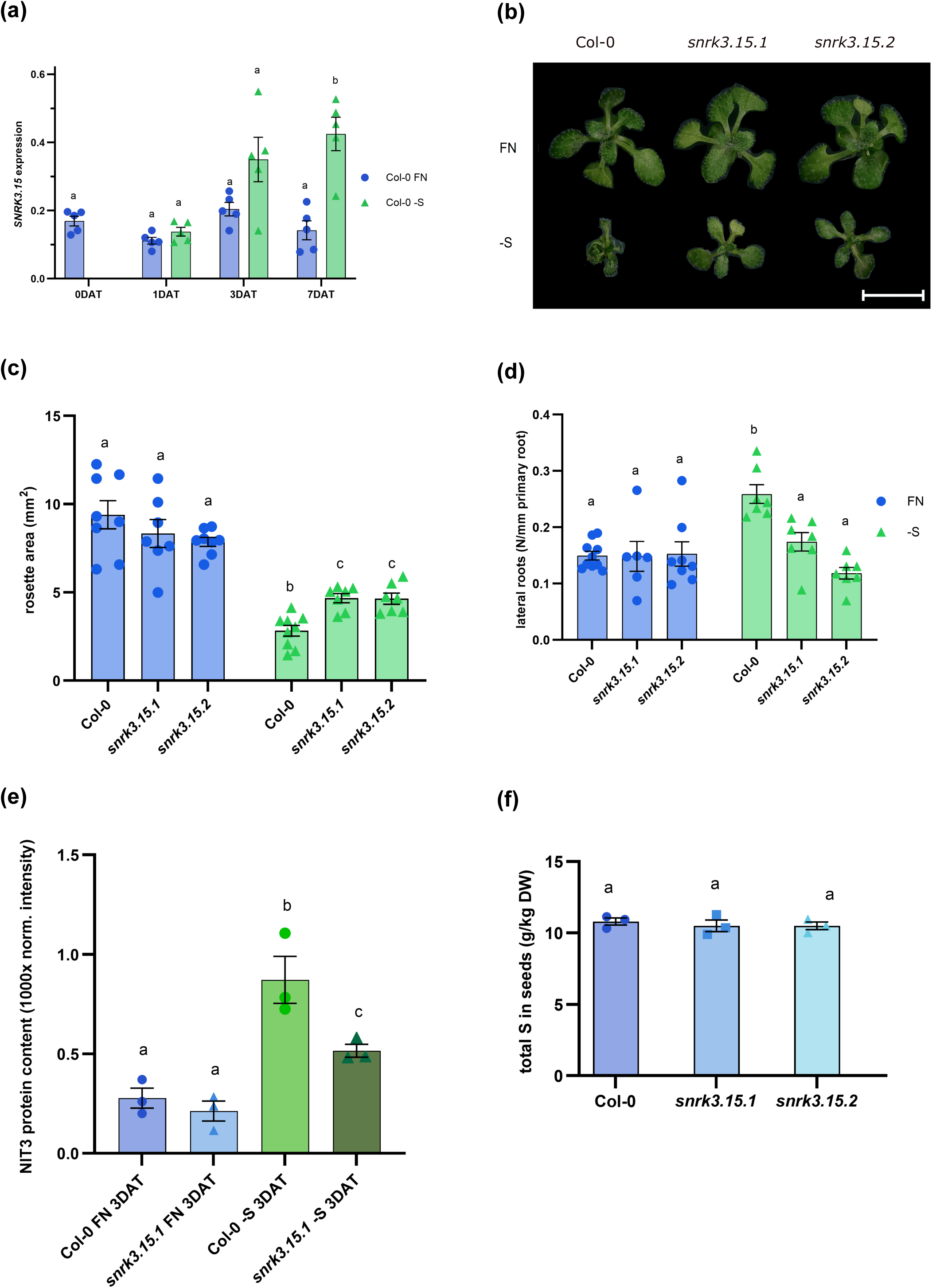
*SNRK3.15* affects growth under S-deficiency. a) *SNRK3.15* transcript levels relative to *TIP41* and *PP2A* determined by qPCR, n = 5. Differences between means were assessed with t-tests between conditions at each timepoint. b) Photo of representative seedling shoots at 16 DAS on plates. Bar length corresponds to 1 cm. c) Rosette area at 16 DAS, n = 7-8 seedlings. (d) Lateral root number, normalized to primary root length, n = 7-8 seedlings. e) NIT3 protein content in microsomal fraction determined by proteomics, n = 3. f) Total sulfur in dry seeds, n = 3. Differences between means were assessed using t-tests (c-f). (a, c – f) Bar height corresponds to the mean, and error bars represent the standard error of the mean. Compact letter display indicates differences with Benjamini-Yekutieli adjusted p ≤ 0.05.

Since *snrk3.15* seedlings grew better than Col-0 under -S (Figure 1b, d) as quantified by determining the leaf rosette area (1c), we investigated whether *snrk3.15* seeds possess more sulfur than Col-0 and thereby provide more sulfur to the growing seedling. ICP-MS was performed, and no difference was found in the concentration of sulfur between Col-0 and *snrk3.15* dry seeds (Figure 1f). To exclude the possibility that *snrk3.15* seeds have more sulfur per seed due to higher seed mass, we determined the seed dry weight of Col-0 and *snrk3.15* mutants and found them to be indistinguishable (Supplemental Figure 2a).

Data for all the elements measured by ICP-MS are presented in Supplemental Figure 2b-e. No differences were noticed in any element in the seeds except for Cu (Supplemental Figure 2d), which was slightly lower in *snrk3.15* seeds than in Col-0. Therefore, despite *snrk3.15* seeds containing an equivalent amount of sulfur as Col-0 seeds, under –S *snrk3.15* mutant seedlings are larger than Col-0.

### Under -S conditions *snrk3.15* mutants have higher chlorophyll contents than Col-0

Sulfur deficiency typically results in decreased chlorophyll (Wulff-Zottele et al., 2010; V. J. Nikiforova et al., 2005), therefore we measured chlorophyll a (Chl-a) and b (Chl-b) contents and calculated the total chlorophyll concentration (Chl-a + Chl-b) (Figure 2a, Supplemental Figure 3a,b). By 7 DAT chlorophyll content was reduced approximately 6-fold in Col-0 and 3-fold in *snrk3.15* relative to FN, resulting in approximately 2-fold higher chlorophyll levels in *snrk3.15* lines than in Col-0 (Figure 2a), which may contribute to the larger rosette phenotype in *snrk3.15* lines (Figure 1c).

**Figure 2:**
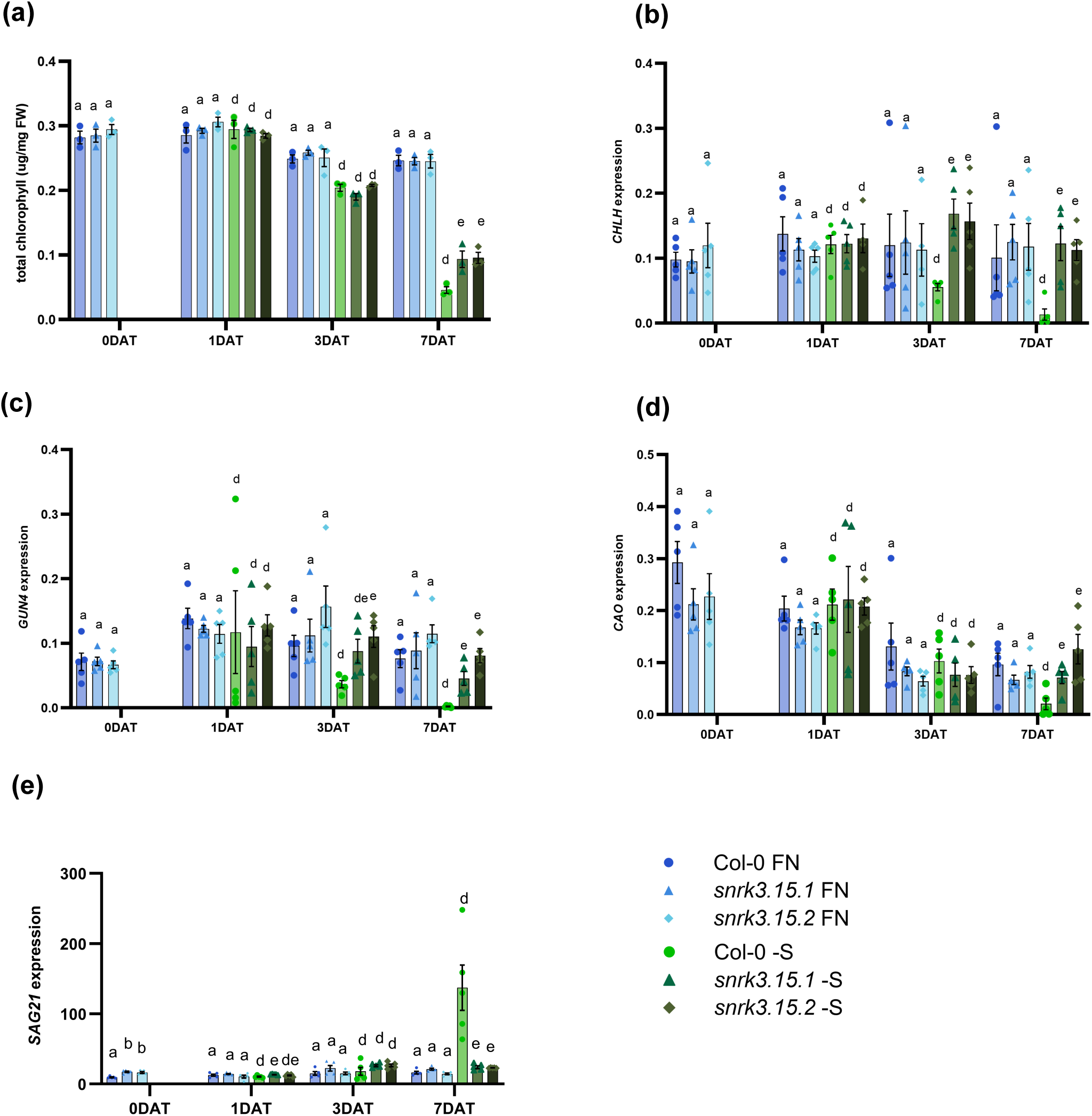
*SNRK3.15* affects chlorophyll response to -S. a) Total chlorophyll content normalized to sample fresh weight, n = 4-5. b – e) Transcript levels relative to *TIP41* and *PP2A* determined by qPCR, n = 5. Bar height corresponds to mean, and error bars represent the standard error of the mean. At each timepoint, differences between means of each *snrk3.15* mutant and Col-0 within each condition were assessed using t-tests. Compact Letter Display indicates differences (Benjamini-Yekutieli adjusted p ≤ 0.05) within each timepoint and condition. Letters a-c are used for FN, and letters d-f are used for -S.

To assess whether the differential decrease in chlorophyll content under -S may be due to differential levels of proteins involved in chlorophyll biosynthesis, proteins were selected based on their annotation to one of the following gene ontology terms: chlorophyll biosynthetic process (GO:0015995), magnesium chelatase complex (GO:0010007), magnesium chelatase activity (GO:0016851), positive regulation of chlorophyll biosynthetic process (GO:1902326), regulation of chlorophyll biosynthetic process (GO:0010380), or protoporphyrinogen IX biosynthetic process (GO:0006782). For each of these proteins detected by LC/MS, the response to -S conditions was determined.

However, none of the 31 proteins showed a differential response to -S in *snrk3.15* compared to Col-0 that was clearly explanatory of the differential chlorophyll content in these lines at 7 DAT (Supplemental Figure 4). Though, even without differential changes in protein abundance, differential abundance at 3 DAT or differential biological activity of those chlorophyll metabolism associated proteins in *snrk3.15* and Col-0 may contribute to the observed differential chlorophyll content.

Since our proteomics data are from 3 DAT, a timepoint before the differential decrease in chlorophyll content was clearly present at 7 DAT, we determined transcript levels of genes involved in chlorophyll biosynthesis throughout the time course. qPCR was performed for three genes with published -S responsiveness (Y. Li et al., 2023; Liu et al., 2020) and at least moderate expression level (Dietzen et al., 2020): CHLH (AT5G13630) is a subunit of magnesium chelatase, which catalyzes formation of Mg protoporphyrin IX, a chlorophyll precursor (W. Zhang et al., 2021; Lichtenthaler, 2012). GUN4 (AT3G59400) is a porphyrin-binding protein responsible for delivering protoporphyrin IX to the magnesium chelatase complex (W. Zhang et al., 2021; Kopečná et al., 2015) and CAO (AT2G47450), which converts chlorophyllide-a to chlorophyllide-b in the last step of chlorophyll biosynthesis (Schumacher et al., 2022). In Col-0 transcripts of *CHLH*, *GUN4*, and *CAO* are strongly downregulated as the duration of sulfate starvation increases (Figure 2b-d), and by 7 DAT, *CHLH*, *GUN4*, and *CAO* expression in Col-0 were 87%, 97%, and 77% lower than at FN, respectively. In contrast, *CHLH* and *CAO* transcript levels in *snrk3.15* mutants were not decreased by the -S condition relative to FN at the same timepoints, a marked difference from their response to -S in Col-0 (Figure 2b,d). GUN4 expression was reduced by –S in *snrk3.15* mutants at 7 DAT but to a lesser extent than in Col-0, resulting in 22.8- fold and 40.7-fold higher transcript levels than in Col-0 (Figure 2c). At 7 DAT to –S, expression of *CAO* was approximately 6.5-fold higher in *snrk3.15* lines than in Col-0 (Figure 2d). At 3 and 7 DAT -S, *CHLH* transcripts were 2.9- and 8.8-fold higher in *snkr3.15.1* and *snrk3.15.2* than in Col-0, respectively (Figure 2b). These data suggest that *SNRK3.15* is necessary for the strong transcriptional downregulation under -S of the chlorophyll biosynthesis genes *CAO*, *CHLH*, and *GUN4*.

The attenuated decrease in chlorophyll content under -S could also be due to differential levels of proteins involved in chlorophyll degradation (Gao et al., 2016; Hörtensteiner, 2006). Therefore, the -S response in Col-0 and *snrk3.15* was determined for the eight proteins in the LC/MS dataset and annotated to the following gene ontology terms: chlorophyll catabolic process (GO:0015996), regulation of chlorophyll catabolic process (GO:0010271), negative regulation of chlorophyll biosynthetic process (GO:1902325). None of the eight proteins showed a response to -S (at 3 DAT) in Col-0 compared to *snrk3.15* that could be clearly associated with lower chlorophyll content at 7 DAT (Supplemental Figure 5). As before, we also performed qPCR for genes important for chlorophyll degradation which have been shown to respond transcriptionally to -S in Arabidopsis seedlings grown on plates (Dietzen et al., 2020): *NYC1* (*AT4G13250*) (Horie et al., 2009) and *SGR1* (*AT4G22920*) (Wang et al., 2023). *NYC1* and *SGR1* transcript levels were similar in Col-0 and *snrk3.15* lines under –S (Supplemental Figure 3c,d). Taking together our chlorophyll metabolism associated transcript and protein abundance data, we suggest that the difference between Col-0 and *snrk3.15* mutants in chlorophyll content at 7 DAT to -S may rather be due to SNKR3.15-regulated activity of chlorophyll metabolism associated proteins.

Chlorophyll degradation is a hallmark of leaf senescence (Gao et al., 2016), and senescence is affected by various parameters including nutrient availability (Whitcomb et al., 2014). Senescence associated gene 21 (*SAG21*) (*AT4G02380*) is often used as an early transcriptional marker of senescence (Ahmad & Guo, 2019; Jing et al., 2002). *SAG21* transcript levels were induced 8.45-fold by –S in Col-0 at 7 DAT but they were not induced appreciably by –S in *snrk3.15* mutants (Figure 2e). The *SAG21* transcript data indicate *snrk3.15* lines having a somewhat delayed senescence onset in -S relative to Col-0, which is consistent with the higher Chl levels in *snrk3.15* lines than in Col-0 in –S at 7 DAT.

### SNRK3.15 is necessary for anthocyanin accumulation under -S conditions

Early leaf senescence is also typically accompanied by accumulation of anthocyanins, a class of flavonoids (Pei et al., 2024). Anthocyanins also accumulate in response to sulfur deficiency (De Kok et al., 2017; V. Nikiforova et al., 2003). Anthocyanin levels in Col-0 were 1.78-fold higher in -S than FN at 7 DAT, while in the *snrk3.15* lines they were not affected by transfer to -S (Figure 3a). The -S response of transcripts for three genes related to anthocyanin levels were assessed, specifically the TFs *MYB75* (AT1G56650), also called *PRODUCTION OF ANTHOCYANIN PIGMENT1* (*PAP1*), and *MYB90/PAP2* (AT4G29080), as well as *ANTHOCYANIDIN SYNTHASE* (*ANS)* (AT4G22880) (Jiang et al., 2022; Bhargava et al., 2010; Zuluaga et al., 2008; Tohge et al., 2005). By 7 DAT the transcripts from all three genes accumulated in –S treated Col-0, while in *snrk3.15* mutants, the induction of their transcript levels was either attenuated (*MYB75/PAP1* and *ANS*) or not observed (*MYB90/PAP2*) (Figure 3b-d).

**Figure 3:**
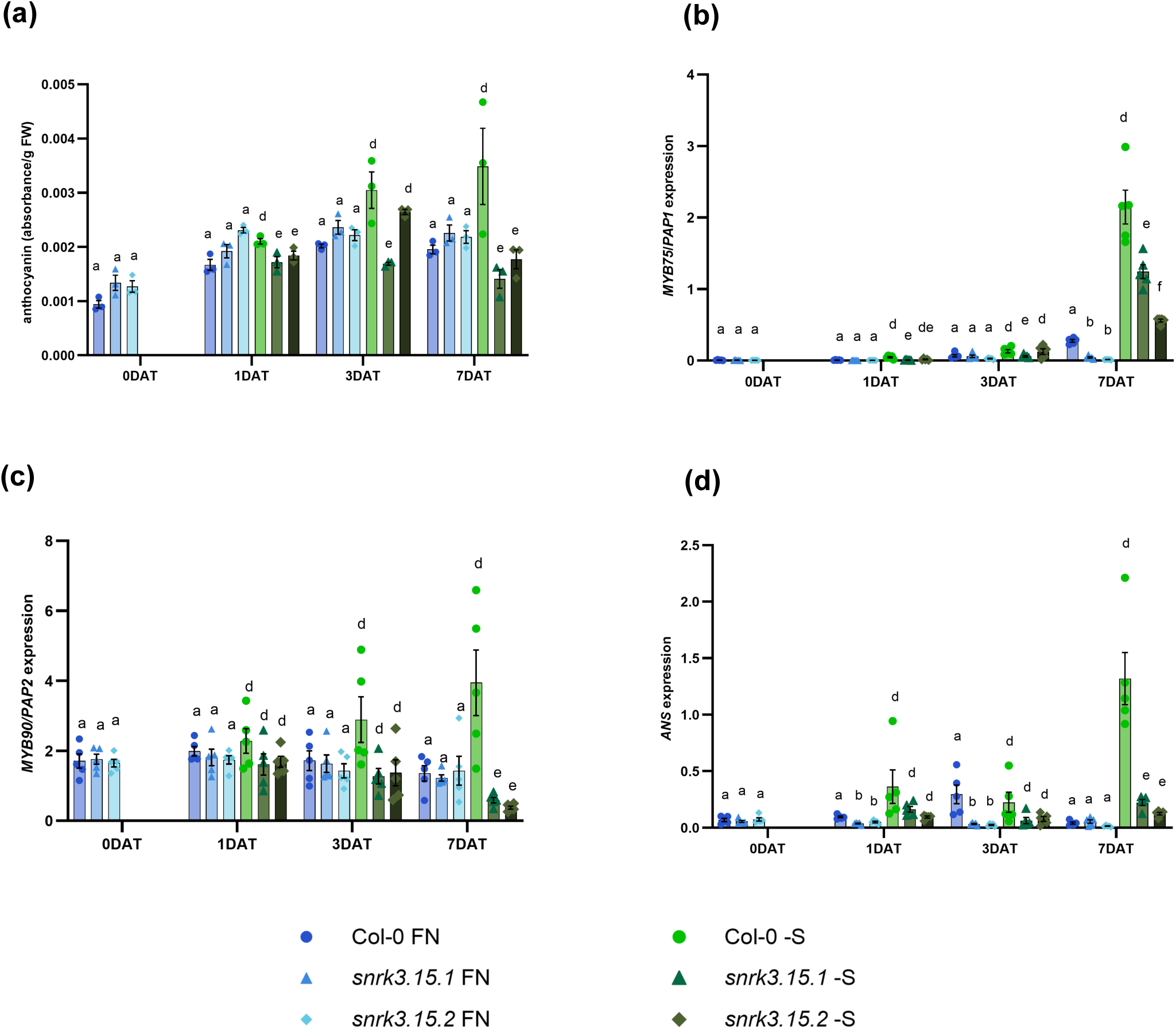
*SNRK3.15* affects anthocyanin response to -S. a) Total anthocyanin content normalized to sample fresh weight, n = 4-5. b – d) Transcript levels relative to *TIP41* and *PP2A* determined by qPCR, n = 5. Bar height corresponds to mean, and error bars represent the standard error of the mean. At each timepoint, differences between means of each *snrk3.15* mutant and Col-0 within each condition were assessed using t-tests. Compact Letter Display indicates differences (Benjamini-Yekutieli adjusted p ≤ 0.05) within each timepoint and condition. Letters a-c are used for FN, and letters d-f are used for -S.

### Changes to sulfur compound content and OAS-cluster gene expression at 1 DAT to -S are attenuated in *snrk3.15*

Transfer of seedlings from FN to -S liquid media resulted in large decreases in internal sulfate concentrations, due to mobilization and metabolism of stored sulfate. At 1 DAT, sulfate was decreased by 78% in Col-0 and by 56% in *snrk3.15* mutants compared to FN, resulting in approximately 2-fold higher sulfate concentration in *snrk3.15* lines than in Col-0 at this early timepoint (Figure 4a). Sulfate concentration continued to decrease in both Col-0 and *snrk3.15* lines as the starvation continued. At 3 and 7 DAT no significant differences in sulfate concentration between *snrk3.15* and Col-0 were found any more.

**Figure 4:**
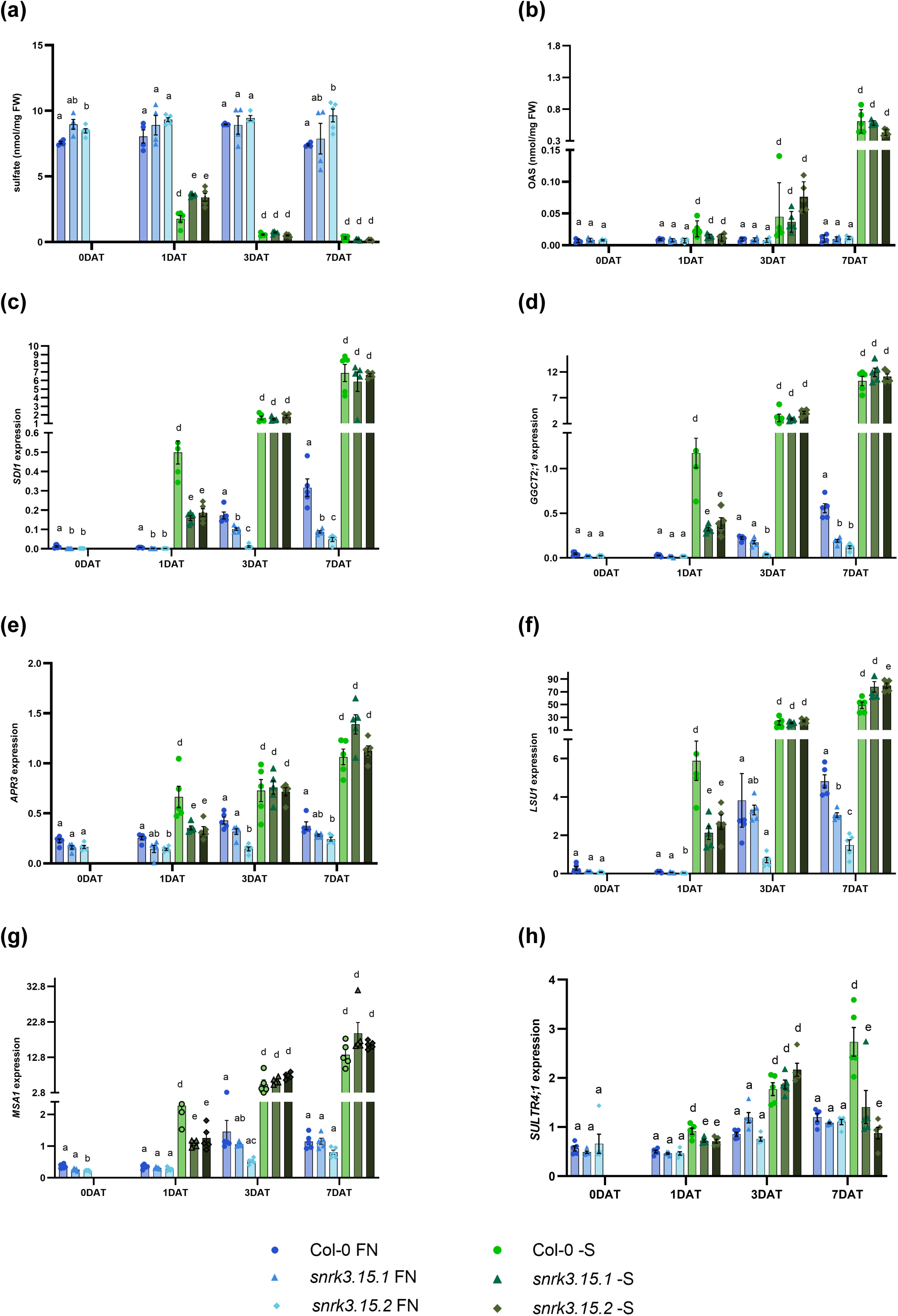
*SNRK3.15* affects early changes to S-deficiency response pathway. a) Sulfate content and b) OAS content normalized to fresh weight, n = 4-5. c – h) Transcript levels relative to *TIP41* and *PP2A* determined by qPCR, n = 5. Bar height corresponds to mean, and error bars represent the standard error of the mean. At each timepoint, differences between means of each *snrk3.15* mutant and Col-0 within each condition were assessed using t-tests. Compact Letter Display indicates differences (Benjamini-Yekutieli adjusted p ≤ 0.05) within each timepoint and condition. Letters a-c are used for FN, and letters d-f are used for -S.

Sulfate deprivation leads to increasing OAS levels during continued starvation, and the concentration of OAS is considered a signaling molecule in S-deficiency responses (Hubberten et al., 2012). At 1 DAT to –S, OAS was approximately 2-fold higher in Col-0 than in *snrk3.15* mutants (Figure 4b), an effect size comparable to that observed for sulfate at this time point (Figure 4a). However, the p-values for these differences in OAS content were slightly higher than 0.05 (0.079 and 0.082 for *snrk3.15.1* and *snrk3.15.2*, respectively). As for sulfate levels, OAS levels at 3 DAT and 7 DAT to -S were comparable in Col-0 and *snrk3.15* lines.

Since OAS has been shown to regulate the expression of OAS-cluster genes (Hubberten et al., 2012), and these genes often serve as transcriptional markers for S deficiency, their transcript levels were determined. In –S, *SDI1, GGCT2;1, APR3, LSU1,* and *MSA1* transcript levels were lower in *snrk3.15* mutants compared to Col-0 at 1 DAT, while as the starvation was extended to 3 and 7 days, their transcript levels were similar in Col-0 and *snrk3.15* mutants (Figure 4c-g), corresponding to the respective sulfate and OAS levels. At FN, *SDI1* is expressed less in *snrk3.15* mutants compared to Col-0 at each timepoint (Figure 4c).

Most sulfate in plant tissue is stored in the vacuole, and since sulfate levels decreased in Col-0 more strongly between 0 DAT and 1 DAT than in *snrk3.15* lines, we determined transcript levels of the main sulfate transporter involved in efflux of sulfate from the vacuole, *SULTR4;1* (De Kok et al., 2017; Kataoka et al., 2004). *SULTR4;1* transcripts increased steadily in Col-0 over the duration of -S treatment, while in *snrk3.15* mutants *SULTR4;1* transcripts increased only until 3 DAT and then returned to FN level by 7 DAT (Figure 4h). At 1 DAT to -S, *SULTR4;1* transcript levels were slightly higher in Col-0 than in *snrk3.15*, which is consistent with higher sulfate efflux from the vacuole resulting in higher utilization in Col-0 during this period.

Based on the data above, we suggest that SNRK3.15 is positively involved in increasing utilization of sulfate in the initial phase of sulfate deprivation, most likely by increasing mobilization of vacuolar sulfate via induction or activation of SULTR4;1. OAS accumulation and induction of the OAS cluster genes strictly follows in an inverse manner the availability of sulfate.

Cys, GSH, and GSL are major pools of organic S-compounds (V. J. Nikiforova et al., 2005). The concentrations of these compounds were reduced after the shift to -S media with increasing reduction with continued starvation (Figure 5). The greater Cys reduction in Col-0 (23%) than in *snrk3.15* mutants (15% and 2%, respectively) at 1 DAT, and higher Cys concentration in *snrk3.15* mutants at 0 DAT than Col-0, resulted in Col-0 containing less Cys than *snrk3.15* at 1 DAT to -S (Figure 5a). GSH, a cysteine containing tripeptide, follows the cysteine concentration pattern, though it is much more strongly reduced under prolonged starvation than Cys (Figure 5b). As with sulfate, OAS, and OAS-cluster gene expression, Cys and GSH show a genotype effect at 1 DAT, but not at later timepoints of S-deficiency. GSLs display a comparable pattern to Cys and GSH over starvation time though with a less significant effect at 1 DAT (Figure 5c).

**Figure 5:**
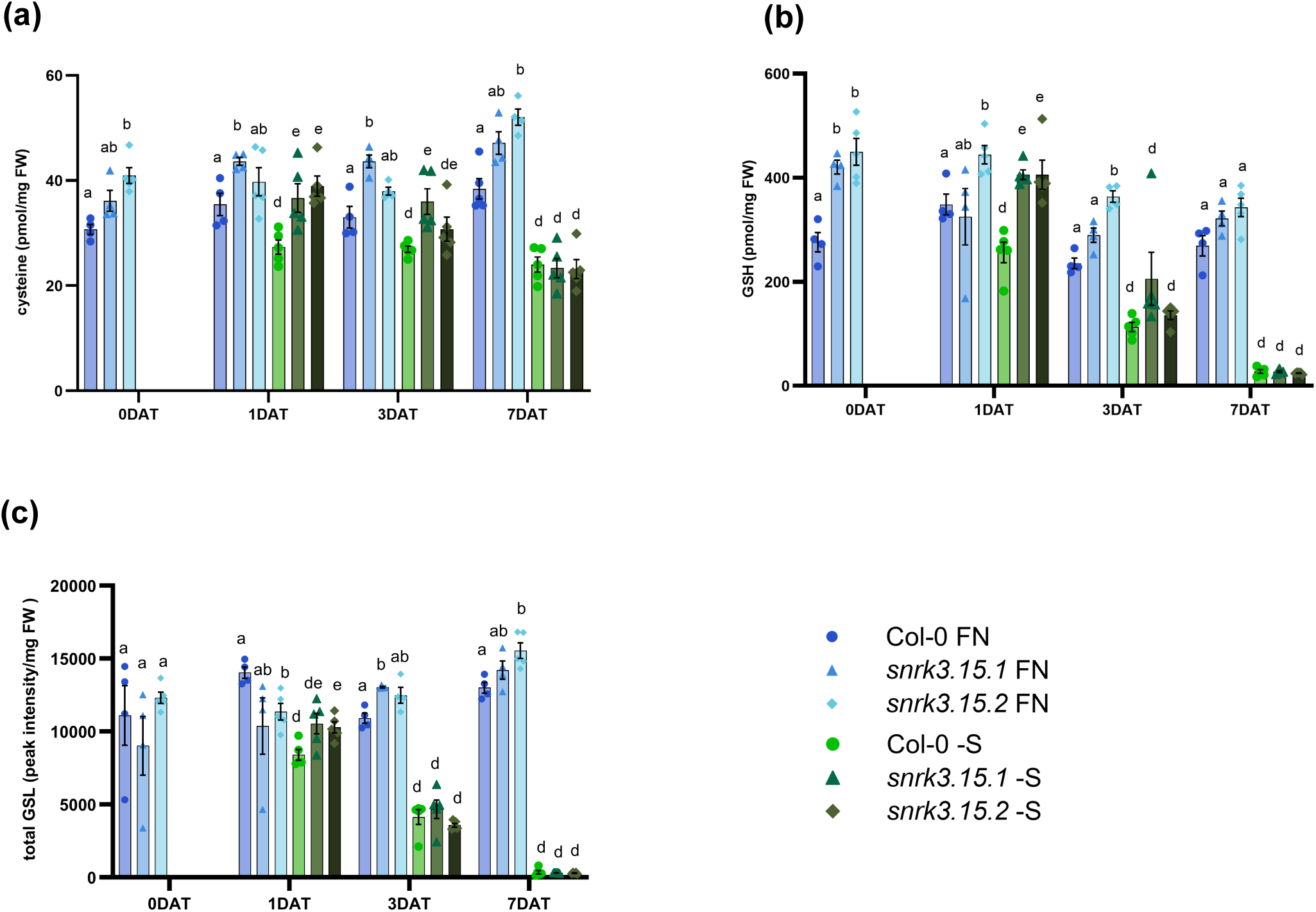
*SNRK3.15* affects early changes to S-metabolite levels in response to -S conditions. a) Cysteine, b) glutathione, and c) total glucosinolate levels normalized to fresh weight, n = 4-5. Bar height corresponds to mean, and error bars represent the standard error of the mean. At each timepoint, differences between means of each *snrk3.15* mutant and Col-0 within each condition were assessed using t-tests. Compact Letter Display indicates differences (Benjamini-Yekutieli adjusted p ≤ 0.05) within each timepoint and condition. Letters a-c are used for FN, and letters d-f are used for -S.

### Proteome response to -S is qualitatively similar but quantitatively attenuated in *snrk3.15* relative to Col-0

To help further explore the molecular underpinnings of the -S associated phenotypes observed in *snrk3.15* lines, we investigated the proteome response to the shift from FN to -S media and how those responses may differ in Col-0 and *snrk3.15.1*. LC/MS-MS based proteome profiling and comparative analysis were performed on seedlings in shaking culture at 3 DAT to -S and to FN media. Principal component analysis allowed assessment of the overall impact of the three experimental variables: condition (FN, -S), genotype (Col-0, *snrk3.15.1*), and protein fraction (microsomal, soluble) on the normalized intensity of identified proteins in each sample (Supplemental Figure 6a). PC1 strongly separates the samples based on protein fraction and accounts for 64% of the total variation among the proteomes. PC2 accounts for 13% of the total variation and separates the samples based on condition. Notably, PC1 and PC2 do not clearly separate between Col-0 and *snrk3.15.1* samples, suggesting that their global proteome responses to -S were similar. This is supported by a scatter plot of log2FC of all proteins detected in Col-0 and *snrk3.15.1*. The effect of shifting *snrk3.15.1* seedlings to -S media was qualitatively similar to that in Col-0 but quantitatively attenuated. By linear regression, the -S response of a given protein in *snrk3.15*.*1* was on average 57% of the response in Col-0 (Figure 6a).

**Figure 6:**
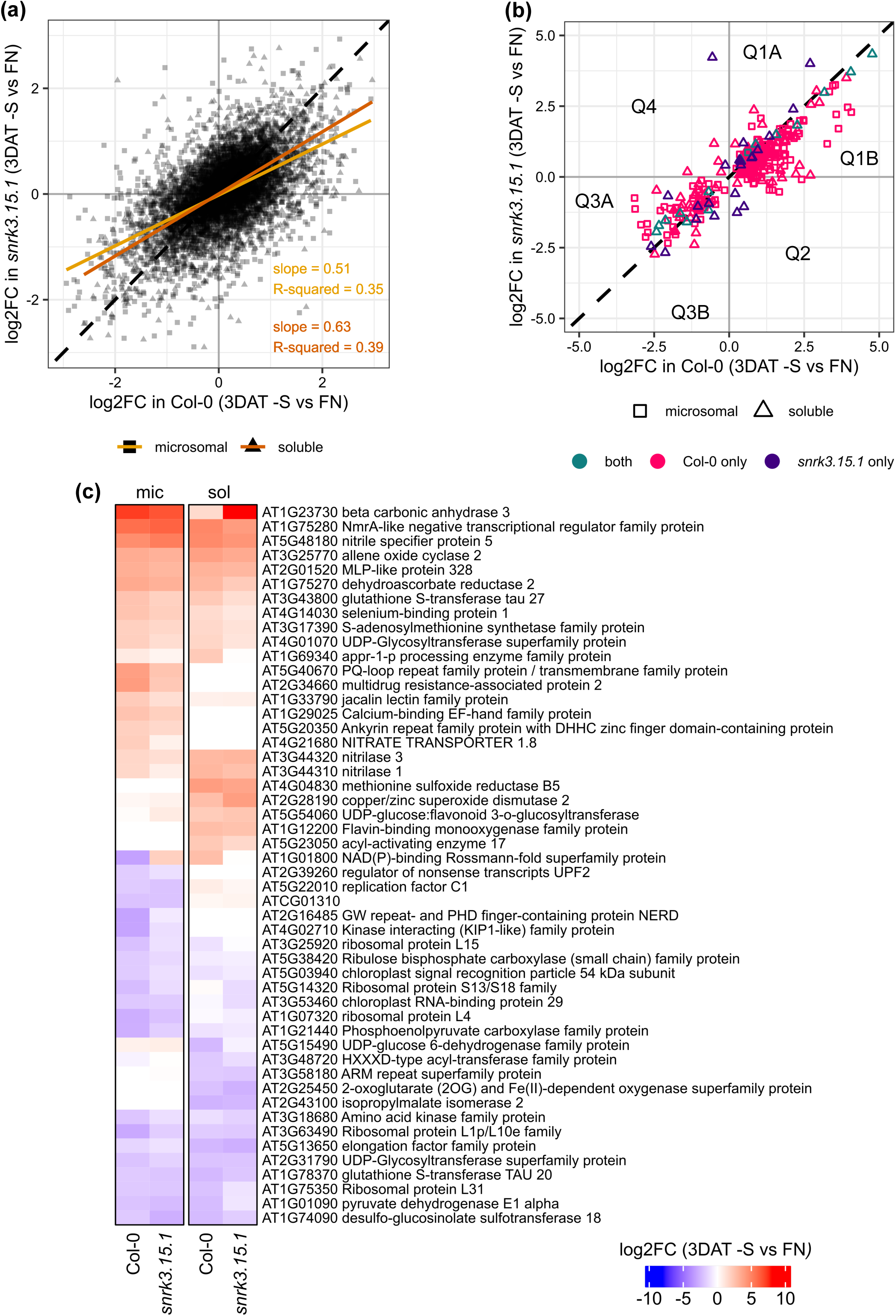
Proteome response to shift to -S media. Col-0 and *snrk3.15.1* shaking cultures were shifted from FN media to either -S or FN media. The proteome response 3 days after transfer (DAT) is represented by the log2FC of each detected protein (a), each DAP (b), and the 25 most strongly increased and decreased DAPs (c). The slope, R-squared value, and line of best fit from the linear regressions for microsomal and soluble samples are drawn on the scatter plot of the global response proteome (a). The DAP subset of the global proteome response is shown in (b), with shape corresponding to proteomic fraction and color indicating whether the protein was classified as a DAP in Col-0 only, *snrk3.15.1* only, or in both Col-0 and *snrk3.15.1*. Quadrants and semi-quadrants of the x-y plane are indicated (Q).

To identify differentially abundant proteins, we performed two types of Welch-tests on microsomal and soluble proteins at 3 DAT: tests for condition-responsive proteins (-S v FN in Col-0 and in *snrk3.15.1*) and for genotype-responsive proteins (*snrk3.15.1* v Col-0 in FN and -S). Proteins with an FDR < 0.1 were classified as differentially abundant proteins (DAP) (Table 1, Supplemental Table 5).

**TABLE 1.**
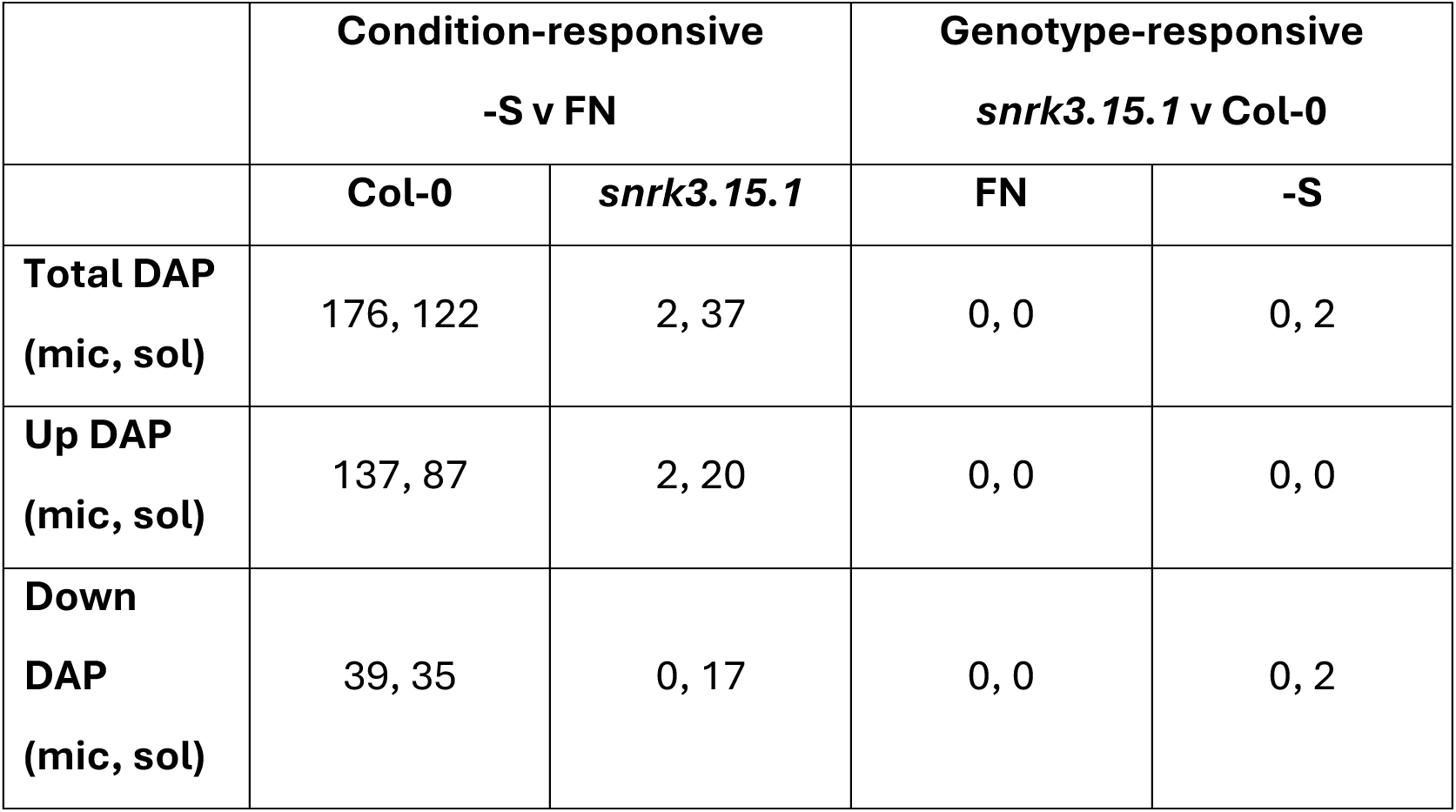

Under sufficient sulfate conditions, *snrk3.15* and Col-0 plants grew very similarly (Figure 1), and no genotype-responsive DAPs were identified in FN seedling samples (Table 1). However, despite the -S associated phenotypes observed in *snrk3.15* at 1 DAT and 7 DAT in shaking culture and on plates (Figure 1-5), only two DAPs between *snrk3.15.1* and Col-0 were identified at 3 DAT to -S: CYSC1/OASTL3;1 (AT3G61440) and PININ (AT1G15200), both of which were less abundant in *snrk3.15.1* than in Col-0 at -S. CYSC1 is catalyzes the formation of ß-cyanoalanine and sulfide in mitochondria from cyanide and cysteine substrates (Hell & Wirtz, 2011). Little is known about PININ function in plants except that it is likely involved in alternative splicing (Bi et al., 2021).

Condition-responsive DAPs were more numerous in both Col-0 and *snrk3.15.1*. In Col-0, 176 and 122 DAPs were identified in the microsomal and soluble protein fractions, respectively, while the corresponding number of DAPs in *snrk3.15.1* was 2 and 37, respectively (Table 1). The attenuated global proteome response to -S in *snrk3.15.1* (Figure 6a) likely accounts for the larger number of DAPs in Col-0. However, the fact that we identified only two microsomal DAPs in *snrk3.15.1* under -S, compared to 176 in Col-0, might rather be due to a localized technical artifact. One of the three biological replicates of the microsomal fraction from *snrk3.15.1* under –S had approximately 40% fewer proteins detected (2228) than the mean proteins detected in all microsomal samples in our dataset (3843) (Supplemental Figure 6b). The normalized intensity values of detected proteins from that biological replicate of the microsomal fraction from *snrk3.15.1* under –S tended to be higher compared to the other two replicates from that group (Supplemental Figure 6c), resulting in higher intragroup variation for proteins in that sample group. Since a larger effect size would be required to achieve the same level of statistical confidence, we suspect that the much smaller number of -S responsive microsomal DAP in *snrk3.15.1* is likely due to a technical artifact, in addition to the identified attenuated proteome response to -S in *snrk3.15.1*.

The differential responses to -S of the DAPs in Col-0 and *snrk3.15.1* are visualized in a scatter plot (Figure 6b). The majority of -S responsive DAPs are found close to the y = x diagonal indicating that the direction of response to -S is the same in Col-0 and *snrk3.15.1*, and that the magnitude of that change is very similar in Col-0 and *snrk3.15.1*. Notably however, most of the 176 microsomal and 154 soluble DAPs were found in the semi-quadrants (Q) 1B or 3A, indicating that for these DAPs, the magnitude of -S response in Col-0 is greater than that in *snrk3.15.1*. Specifically, 104 microsomal and 58 soluble proteins are found in Q1B, and 27 microsomal and 24 soluble proteins are found in Q3A.

The DAPs most strongly affected by transfer to -S were selected, and the log2FCs relative to FN are presented in a heatmap (Figure 6c). Broadly, the magnitude of the response to -S of these most strongly affected proteins is very similar in Col-0 and *snrk3.15.1*. AT1G23730 BETA CARBONIC ANHYDRASE 3 (BCA3) and AT5G48180 NITRILE SPECIFIER PROTEIN 5 (NSP5) are among the most strongly accumulated proteins under -S. BCA3, one of the 6 ß-carbonic anhydrases in Arabidopsis, catalyzes the interconversion of CO_2_ and HCO_3_^−^. ß-carbonic anhydrases are a major constituent of the leaf proteome in Arabidopsis, but their physiological roles are ambiguous (DiMario et al., 2017). However, BCA3 has been recently annotated to sulfur compound metabolic process (GO:0006790) using a multi-omics network-based function prediction method (Depuydt & Vandepoele, 2021). NSP5, participates in GSL catabolism by diverting myrosinase-catalyzed degradation products from isothiocyanate or epithionitrile to nitrile formation (Kong et al., 2012). During GSL catabolism, nitrile and sulfate are released providing N and S sources for other metabolic processes (Falk et al., 2007). Furthermore, the DAP most strongly reduced by transfer to -S media is AT1G74090 DESULFO-GLUCOSINOLATE SULFOTRANSFERASE 18 (SOT18), a protein involved in methionine-derived aliphatic GSL core structure biosynthesis (Harun et al., 2021). *SOT18* transcripts have also been shown to be downregulated under –S conditions in Col-0 (Aarabi et al., 2016). Changes to NSP5 and SOT18 abundance may contribute to the very strong reduction in GSL content under –S conditions (Figure 5c).

Another of the most strongly reduced DAPs is AT1G78370 GLUTATHIONE S-TRANSFERASE TAU 20 (GSTU20), which is involved in cellular detoxification during oxidative stress (Allocati et al., 2018). GSTU20 protein has been previously shown to be strongly reduced in Arabidopsis rosettes by long-term S-starvation (Luo et al., 2021), and glutathione transferase proteins decrease in oilseed rape leaves (D’Hooghe et al., 2013). However, another glutathione S-transferase in the Tau family, GSTU27 (AT3G43800) responds to -S in the opposite direction in our system. While GSTU20 was approximately 4- fold decreased at 3 DAT to -S relative to FN, GST27 was approximately 4-fold increased (Figure 6c).

Since nutrient depletion results in reactive oxygen species (ROS) production (Chandra & Pandey, 2014), we searched in the proteomics data for relevant proteins. Among DAPs that accumulate under -S were seven proteins annotated to cellular oxidant detoxification (GO:0098869), 29 to oxidoreductase activity (GO:0016491), and eight to response to oxidative stress (GO:0006979) GO terms. Out of the 29 DAPs with oxidoreductase activity, five were among the 25 most strongly accumulated proteins under –S: NmrA-like (AT1G75280), DHAR2 (AT1G75270), MSRB5 (AT4G04830), CSD2 (AT2G28190), and FMO (AT1G12200) (Figure 6c). NmrA-like, DHAR2, and MSRB5 have also been found to be highly accumulated in sulfur-starved Arabidopsis rosettes (Luo et al., 2021).

Even though the proteome response to -S was very similar in *snrk3.15.1* and in Col- 0, albeit attenuated overall in *snrk3.15.1*, 22 of 39 condition-responsive DAPs in *snrk3.15.1* were not found to be differentially abundant in Col-0 (Table 2). Six of the 22 -S responsive *snrk3.15*-specific DAPs are predicted to have oxidoreductase activity (GO:0016491), including CSD2 (copper/zinc superoxide dismutase 2, AT2G28190), whose content was 16- fold higher under –S than in FN in *snrk3.15.1*, but only 6.5-fold higher in Col-0. CSD2 degrades reactive oxygen species, specifically superoxide radicals, which accumulate in plants under –S (Chandra & Pandey, 2014). Having higher CSD2 protein levels may contribute to better mitigation of ROS generation in *snrk3.15* seedlings than in Col-0.

**TABLE 2.**
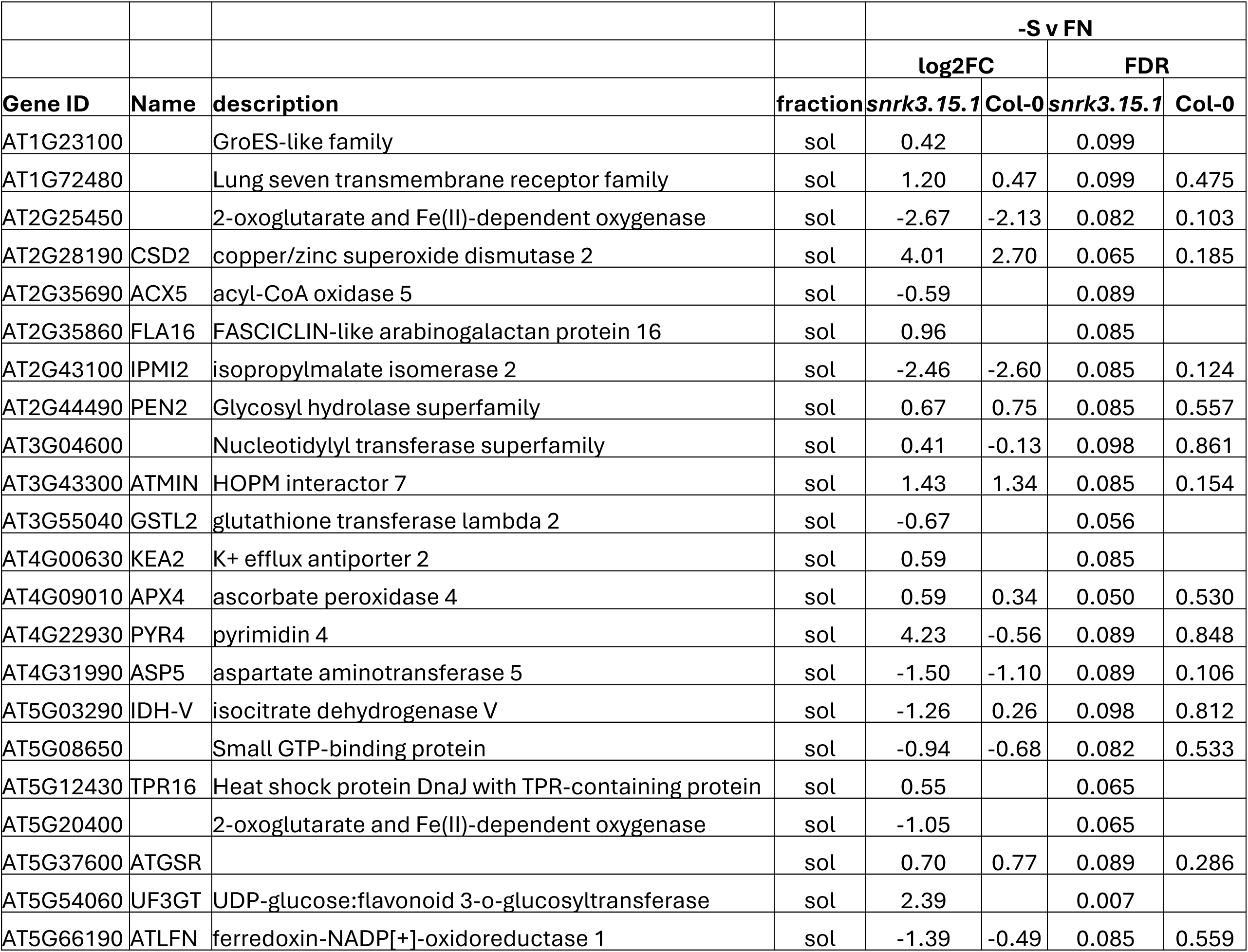

Inspection of the normalized intensity values for these 22 *snrk3.15*-specific DAPs (Supplemental Figure 7) reveals that most did not meet the DAP criteria in Col-0 due to lack of detection in a Col-0 sample or high intra-group variance in Col-0, not due to lack of response to -S conditions.

## DISCUSSION

Sulfate depletion results in numerous metabolic, molecular, and phenotypic alterations in Arabidopsis (Aarabi et al., 2020; Watanabe et al., 2010, 2012). These alterations allow the plant to respond to low sulfate availability through a series of interlaced adaptation processes including increasing uptake capacities, reductive assimilation rates, and remobilization of internal reserves such as vacuolar sulfate and breakdown of S-rich biomolecules such as GSL and GSH. Growth rates decline, and if these coping mechanisms are unable to restore homeostatic S-levels, the plant increases autophagy, enters senescence, and sets seeds prematurely. In this work, we have shown that SNRK3.15 is involved in both the early adaptive responses and the later emergency salvage processes of the annual plant, Arabidopsis.

Kinases are often key players in transducing nutrient status signals to molecular components involved in metabolic and developmental program regulation. Despite the physiological importance of sulfur, to date no signaling kinases have been identified in sulfur-deficiency signaling response programs. *SNRK3.15* expression is upregulated by shifting seedlings to -S (Figure 1a), and it belongs to the SNRK3 kinase family, several members of which have been shown to be involved in deficiency responses to other nutrients. Further, *SNRK3.15* expression is regulated by a key S-response TF, SLIM1. Therefore, we suspected that SNRK3.15 may participate in Arabidopsis S-deficiency responses.

Indeed, when Arabidopsis Col-0 seedlings were exposed to sulfate starvation they demonstrated previously described –S responses including reduced growth (Figure 1b,c), altered root architecture (Figure 1d), and changes at the molecular level including reduction in levels of chlorophyll (Figure 2a, Supplemental Figure 3a,b), sulfate (Figure 4a), cysteine, GSH, and GSL (Figure 5), accumulation of OAS and induction of OAS-cluster genes (Figure 4b-h), and a significant shift in the proteome (Figure 6, Table 1, Table 2). In contrast, *snrk3.15* mutants displayed these responses to -S in a delayed or attenuated manner, particularly at the early stages of the treatment, as if the mutants did not fully sense the sulfate deprivation or did not fully transduce the respective signals. The general attenuation in S-deficiency responses in *snrk3.15.1* and *snrk3.15.2* relative to Col-0 indicates that SNRK3.15 kinase contributes to establishing molecular and physiological responses to sulfate deprivation.

Among the early responses to shifting plants to -S are changes in sulfate, OAS, and OAS-cluster gene expression, all of which showed attenuated responses in *snrk3.15* relative to Col-0. Notably, the negative relationship between sulfate and OAS and between sulfate and OAS-cluster gene expression is very similar in Col-0 and in *snrk3.15* (Supplemental Figure 8). This suggests that shortly after the shift from FN to -S, SNRK3.15 regulates internal sulfate levels but not downstream OAS accumulation and OAS-cluster transcript induction. SNRK3 kinases have previously been shown to regulate nitrate and phosphate transporter activity, therefore, we hypothesize that SNRK3.15 may regulate vacuolar SULTR activity as an early response to the shift from FN to -S media, resulting in lower sulfate efflux from the vacuole and higher total internal sulfate in *snrk3.15* mutants. It may be that *snrk3.15* mutants are initially partially deficient in sensing or properly transducing the lack of sulfate in the growth media or initial decreases in internal sulfate content. Further, *snrk3.15* seedlings did not increase lateral root density as a response to insufficient S as was observed for Col-0 in this experimental setup. This supports a role for SNRK3.15 in sensing internal sulfur resources and/or a role in transducing that signal.

Interestingly, from 3 DAT many of the early -S response phenotypes, including sulfate, Cys, and GSH accumulation, had equalized between Col-0 and the *snrk3.15* mutants. However, *snrk3.15* seedlings were approximately 70% larger than Col-0 seedlings at 11 DAT, as measured by rosette area (Figure 1c). We tested whether this seedling growth advantage of *snrk3.15* mutants on -S media may be due to higher S content in the seeds, which could have accumulated during prior growth on sufficient sulfate supply. As this is not the case (Figure 1f), we reasoned that SNRK3.15 is not critical for S-content control in seeds under sufficient sulfur conditions, but it is involved in downstream regulatory processes that affect plant growth under S-starvation. Analysis of the metabolome and transcriptome in future studies may shed more light on the causes of the observed growth advantage of *snrk3.15* mutants under S-deprivation.

Prolonged S-deficiencies lead to nutrient depletion induced senescence (NuDIS), which is characterized by decreased chlorophyll content, protein and RNA degradation, and reduced growth (Watanabe et al., 2010). At 7 DAT, the senescence marker gene *SAG21* was clearly induced by –S in Col-0 seedlings, but not in *snrk3.15.1* or *snrk3.15.2* mutants (Figure 2e). This suggests that at 7 DAT, Col-0 seedlings have already entered a NuDIS program, but *snrk3.15* mutants are delayed in NuDIS entry. Differential decrease in chlorophyll content supports this conclusion (Figure 2a, Supplemental Figure 3a,b).

Chlorophyll content decreases in both Col-0 and *snrk3.15* mutants upon extended S-deficiency, but at 7 DAT, the reduction is attenuated in both *snrk3.15* mutants relative to Col-0. Transcript levels of several key chlorophyll biosynthesis genes (*GUN4*, *CAO*, *CHLH*) are stronger down-regulated in Col-0 seedlings than in *snrk3.15* mutants at 3 and 7 DAT to -S. The levels of many proteins that positively affect chlorophyll content were determined, and some showed strong reductions under -S, including CHLH. However, the levels of most were either not clearly affected by -S at 3 DAT, or the -S response was similar in Col-0 and *snrk3.15.1*, as was the case for CHLH (Supplemental Figure 4). With the data presented here, overall, it is hard to identify a mechanism for the observed attenuated chlorophyll reduction in *snrk3.15*, but the data do clearly support SNRK3.15 involvement in triggering NuDIS traits.

Increasing levels of the phytohormone ABA is important for senescence developmental programs such as NuDIS. However, ABA levels tend to decrease under S-starvation, perhaps because Cys, which positively affects ABA biosynthesis, is strongly decreased under sulfate depletion (Cao et al., 2014). SNRK3.15 is a central hub controlling ABA-responsive genes (Lumba et al., 2014), which may allow recruitment of ABA-dependent responses under sulfate deprivation such as NuDIS. Some of the observed attenuated or delayed responses to S-starvation in the *snrk3.15* mutants may be due to weaker activation of ABA-dependent senescence pathways.

Open questions remain regarding SNRK3.15 regulation and function. SNRK3 family members (26 in Arabidopsis) require interaction with calcium binding proteins, specifically CBL family proteins (10 in Arabidopsis), for kinase activity, but CBL-SNRK3 pairings are typically non-exclusive. SNRK3.15 has been shown to interact *in planta* with CBL2, CBL3, and CBL8, resulting in distinct CBL-SNRK3.15 complexes in different subcellular compartments (Batistič et al., 2009). However, in the context of S-deficiency the localization and primary upstream CBL regulators of SNRK3.15 activity are unknown. What are the targets of SNRK3.15 kinase activity during early responses and later responses to S-deficiency, and how does phosphorylation of those targets affect their activity and contribute to S-deficiency responses including NuDIS? More broadly, what are the components and structure of the signal transduction cascades relevant to S-deficiency responses and where does SNRK3.15 kinase fit into these signal transduction pathways?

Knowledge of the molecular mechanisms and signaling regulating sulfur deficiency lag that for two other critical macronutrients, nitrogen and phosphorus (Q. Li et al., 2020; Ristova & Kopriva, 2022) . In summary, we speculate that SNRK3.15 is an upstream regulator of various stress response pathways. As a SLIM1-dependent regulatory factor, it responds early to decreases in sulfate content in plant tissues. SNRK3.15 is likely involved in ramping down metabolism, allowing the plant to endure phases of sulfate deprivation by sparing resources, reducing potentially dangerous sources of ROS by reducing chlorophyl content and hence photosynthesis, and eventually slowing growth. When sulfate depletion leads to severe effects on biosynthetic capacity and physiology, plants enter senescence. Given that under sulfate depletion ABA biosynthesis is impaired, SNRK3.15 might be a crucial factor for the plant to enter NuDIS, which hence allows the plant to run an emergency program with earlier flowering and shift remaining resources into seed production. With the onset of this NuDIS program we assume that resupplying sulfate will probably not alleviate starvation symptoms or reverse the senescence program. Whether an effect of SNRK3.15 on the TOR/SNRK1 system exists can at this point only be speculated, but given the very broad pleiotropic effects it appears likely.

## ACKNOWLEDGMENTS

AA was funded by the Deutsche Forschungsgemeinschaft (DFG), HO1916/13–1. SM and SK are funded by the DFG under Germany’s Excellence Strategy – EXC 2048/1 – project 390686111 and project 426501900 (SK). RH, PP, AA, WS, SA, EH, and SJW received funding from the Max Planck Society. The authors would like to especially thank Franziska Brueckner for performing ion chromatography and helping with processing the huge number of samples and Sabine Ambrosius for assistance with the elemental analysis. Many thanks to Arun Sampathkumar for mentoring AA regarding the microscope handling in order to produce the seedling figures presented in the manuscript.

## AUTHOR CONTRIBUTIONS

AA designed and performed the experimental work, analyzed the data, and prepared the graphs. EH confirmed the zygosity of the SALK T-DNA lines SALK_009699 *(snrk3.15.1*) and SALK_147899 *(snrk3.15.2*) and prepared the samples for proteomics. WS performed the proteomic profiling. SA performed LCMS and selected the peaks from the chromatograms. PP performed the cysteine chromatography and selected the peaks. SM and SK were responsible for the seed element analysis. RH managed the funding acquisition, coordinated the project, and supervised AA. SJW analyzed the proteomic data and supervised AA. AA, RH, and SW wrote and edited the manuscript.

