## Supplemental Figures 1-8 for "*SNRK3.15* is a crucial component of the sulfur deprivation response in *Arabidopsis thaliana*"

#### Supplemental Figure Captions

##### Supplemental Figure 1: Supplemental characterization of *snrk3.15* lines, *SNRK3.15* expression, and comparative growth in soil

a) Scheme of the Arabidopsis *SNRK3.15* gene. Exon is indicated as a solid black box. The position of the T-DNA insertion is shown for both T-DNA mutations (SALK_09699 and SALK_147899), while the arrows represent the orientation of the insertion. b) *SNRK3.15* transcript level relative to *TIP41* and *PP2A* determined by qPCR. c) Photo of representative Col-0 and *snrk3.15* mutants rosettes at 23 DAS grown in the greenhouse under long day conditions (16h day - 8h night). d) Rosette area of Col-0 and *snrk3.15* mutants, grown under long day conditions (16h day - 8h night) in the greenhouse. b, d) Bar height corresponds to mean of 4-5 replicates, and error bars represent the standard error of the mean. At each timepoint, statistical significance was assessed using t-tests between *snrk3.15* and Col-0. Compact letter display identifies *snrk3.15* mutants that are statistically different than Col-0 (Benjamini-Yekutieli adjusted $p \leq 0.05$). Letters a-c were used for comparisons at FN. Letters d-f were used for comparisons at -S.

##### Supplemental Figure 2: Phenotypes of dry seed from Col-0 and *snrk3.15* plants grown on soil

Mature seeds were collected from siliques and dried in a drying room with humidity maintained at 15% and temperature at 15°C. a) Mean single seed weight was calculated from the weight of 50 seeds. Bar height corresponds to mean calculated single seed weight from the 5 independent seed pools, and error bars represent the standard error of the mean. Compact letter display identifies *snrk3.15* mutants that are statistically different than Col-0 (t-test, Benjamini-Yekutieli adjusted $p \leq 0.05$). b-e) Elemental profile of dry seeds, as determined by inductively coupled plasma mass spectrometry. Bar height corresponds to mean of 3 replicates, and error bars represent the standard error of the mean. For each element, statistical significance was assessed using t-tests between *snrk3.15* and Col-0. Compact letter display identifies genotypes of *snrk3.15* that are statistically different than Col-0 (Benjamini-Yekutieli adjusted $p \leq 0.05$).

##### Supplemental Figure 3: Response of chlorophyll and chlorophyll degradation genes to -S in Col-0 and *snrk3.15* seedlings

a) Chlorophyll-a and (b) chlorophyll-b content in Col-0 and *snrk3.15* mutants. Transcript level of chlorophyll degradation genes, (c) *NYC1* and (d) *SGR1* relative to *TIP41* and *PP2A* determined by qPCR. Bar height corresponds to mean of 4-5 replicates, and error bars represent the standard error of the mean. At each timepoint, statistical significance was assessed using t-tests between *snrk3.15* and Col-0. Compact letter display identifies *snrk3.15* mutants that are statistically different than Col-0 (Benjamini-

Yekutieli adjusted p ≤ 0.05). Letters a-c were used for comparisons at FN. Letters d-f were used for comparisons at -S.

**Supplemental Figure 4: Levels of proteins positively associated with chlorophyll content**

The normalized signal intensity of selected proteins in the proteomics dataset is shown. Each point corresponds to a biological replicate. Point shape corresponds to protein fraction, point fill corresponds to condition (3 DAT to -S or 3 DAT to FN), and point color corresponds to genotype. Proteins potentially positively associated with chlorophyll levels were selected based on their having been annotated to at least one of the following gene ontology terms: chlorophyll biosynthetic process (GO:0015995), magnesium chelatase complex (GO:0010007), magnesium chelatase activity (GO:0016851), positive regulation of chlorophyll biosynthetic process (GO:1902326), regulation of chlorophyll biosynthetic process (GO:0010380), protoporphyrinogen IX biosynthetic process (GO:0006782).

**Supplemental Figure 5: Levels of proteins negatively associated with chlorophyll content**

The normalized signal intensity of selected proteins in the proteomics dataset is shown. Each point corresponds to a biological replicate. Point shape corresponds to protein fraction, point fill corresponds to condition (3 DAT to -S or 3 DAT to FN), and point color corresponds to genotype. Proteins potentially negatively associated with chlorophyll levels were selected based on their annotation to at least one of the following gene ontology terms: chlorophyll catabolic process (GO:0015996), regulation of chlorophyll catabolic process (GO:0010271), negative regulation of chlorophyll biosynthetic process (GO:1902325).

**Supplemental Figure 6: Exploratory analysis of global proteome**

a) PCA scores plot illustrating the distribution of samples in the first two principal component space (PC1 and PC2). The percent of total variance explained by each component is shown on the relevant axis in parentheses. Each point corresponds to a sample. Point shape represents protein fraction, point fill corresponds to condition (3 DAT to -S or 3 DAT to FN), and point color corresponds to genotype. PCA by singular value decomposition was performed on pareto scaled, imputed normalized intensity data matrix. b) histograms of the number of proteins detected in each sample. Samples from microsomal and soluble protein fractions are shown separately. Far fewer proteins were detected in the microsomal fraction sample *snrk3.15*_-S_rep2, as indicated. c) density plots of the normalized intensity values (log10 transformed) of the proteins quantified in at least 6 of 12 samples. Samples from microsomal and soluble protein fractions are shown separately. The sample *snrk3.15*_-S_rep2 has a density trace that is shifted relative to the other 11 microsomal fraction samples, as indicated.

**Supplemental Figure 7: Levels of -S responsive, snrk3.15-specific DAPs**

The normalized signal intensity of those proteins that were found to be differentially abundant in 3 DAT to -S compared to 3 DAT to FN in *snrk3.15.1* but not in Col-0 is shown. Only soluble fraction data are shown, because the 22 proteins that met the snrk3.15-specific and -S responsive criteria did so in the soluble fraction, but not the microsomal fraction samples. No proteins identified in the microsomal faction met the snrk3.15-specific and -S responsive criteria. Each point corresponds to a biological replicate. Point fill corresponds to condition, and point color corresponds to genotype.

**Supplemental Figure 8: Relationship between sulfate, OAS, and OAS-cluster genes**

Scatter plots of OAS concentration (a) or OAS-cluster gene expression (b-f) in relation to sulfate concentration. All data were log10 transformed prior to plotting. Each point corresponds to a single biological sample. Point color corresponds to the genotypic line of the sample. Point shape corresponds to the sample timepoint, days after transfer (DAT). And point fill corresponds to the condition after transfer.

### Supplemental Figure 1

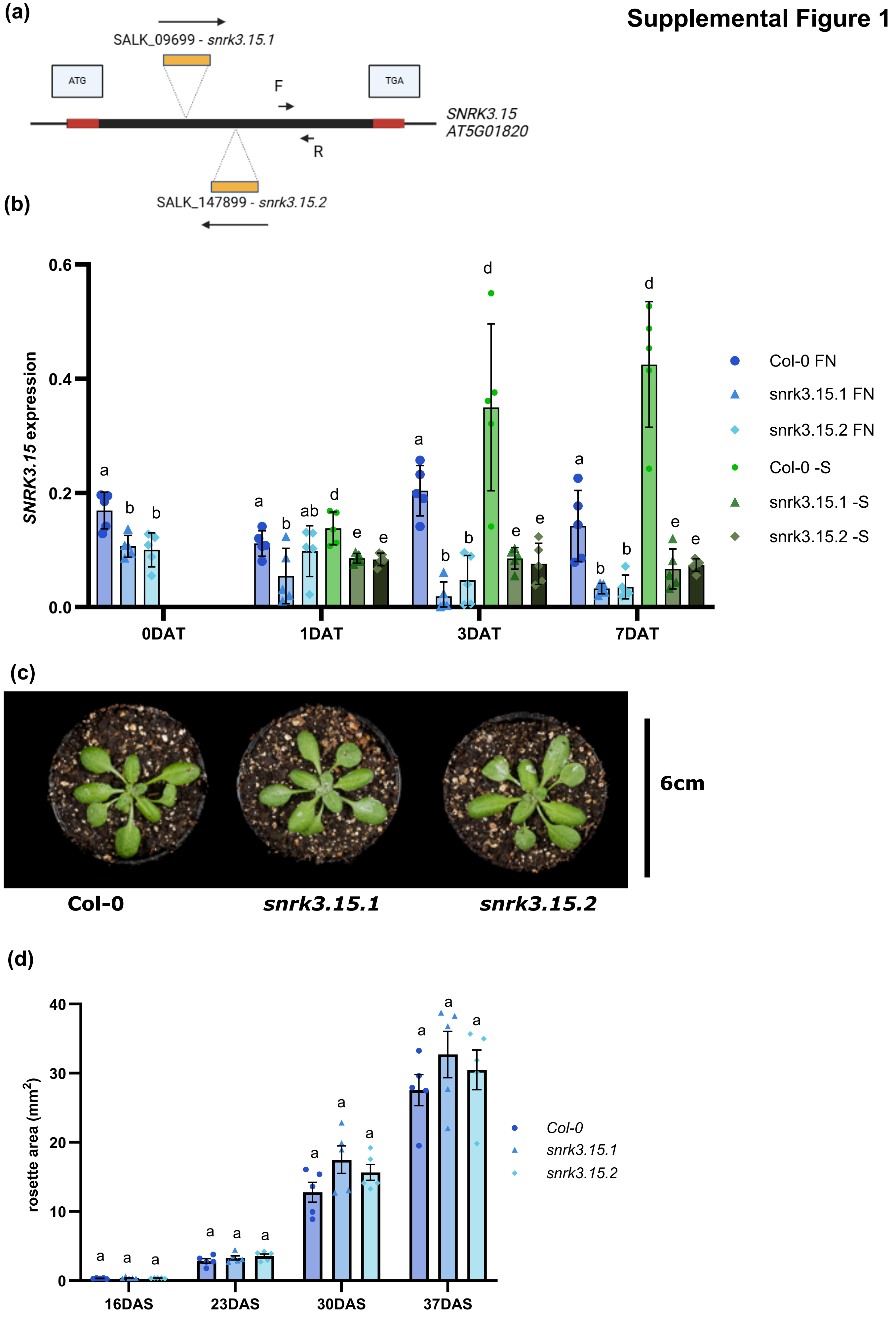

#### Supplemental Figure 2

(a)

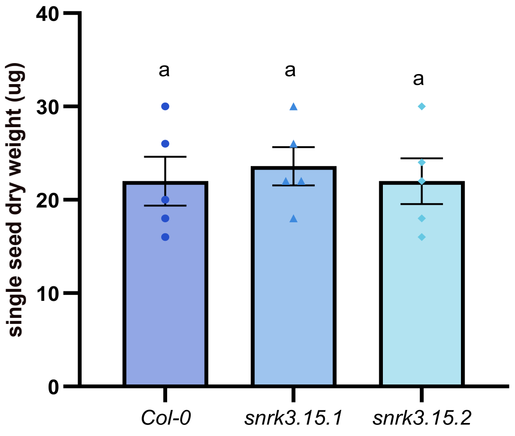


(b)

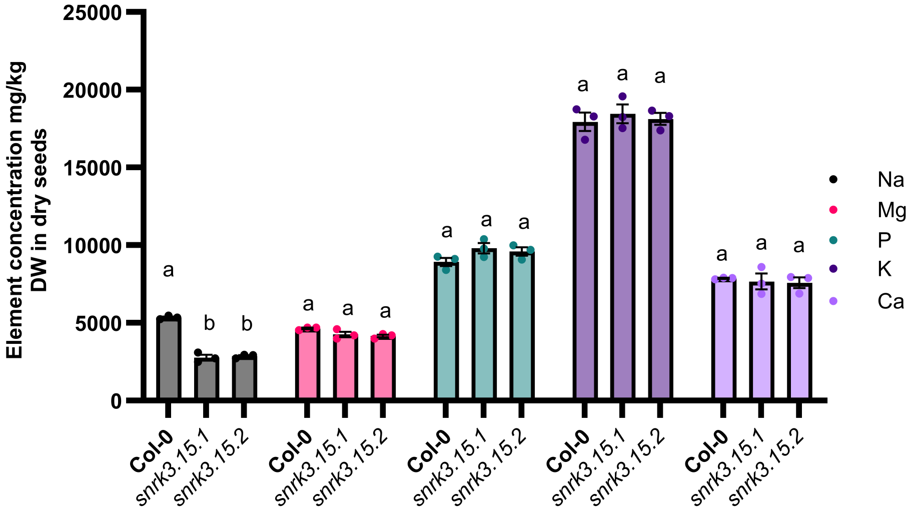


(c)

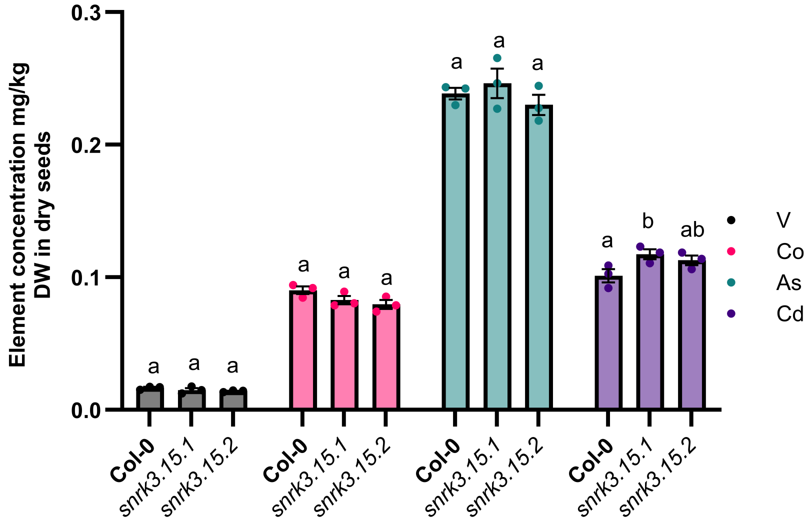

**Supplemental Figure 2**

(d)

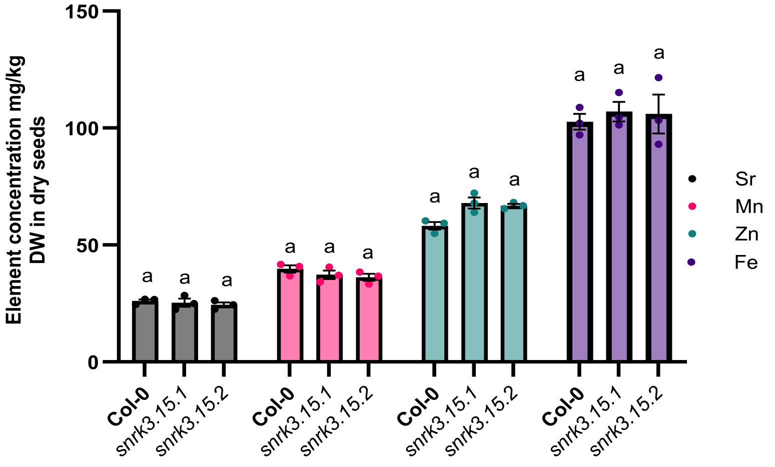


(e)

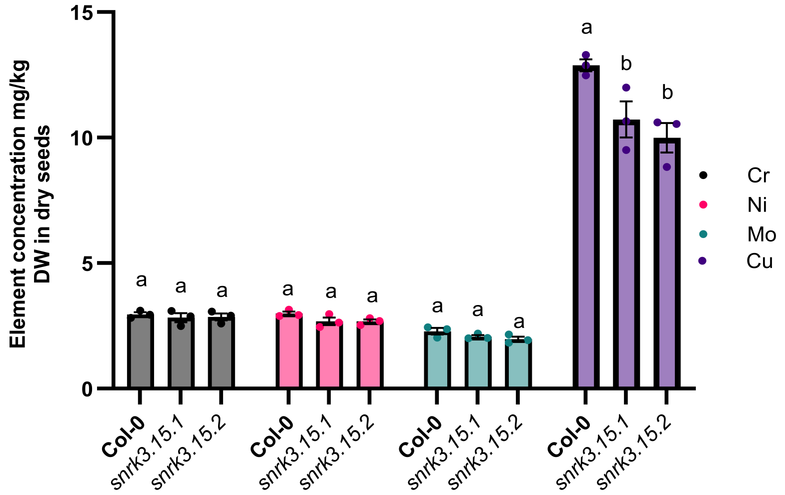

### Supplemental Figure 3

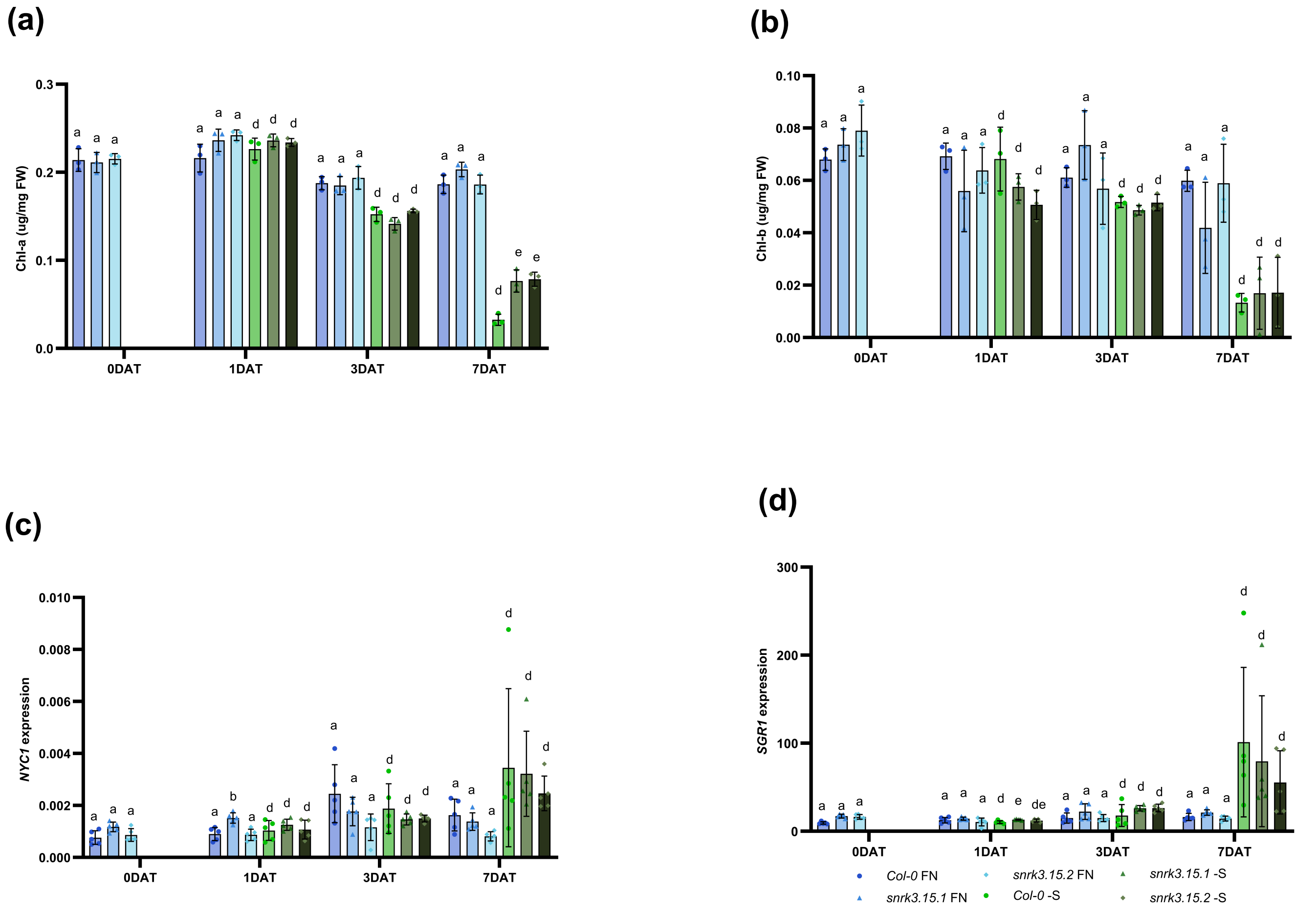

### Supplemental Figure 4

proteins annotated to GO terms positively associated with chlorophyll levels
page 1 of 3

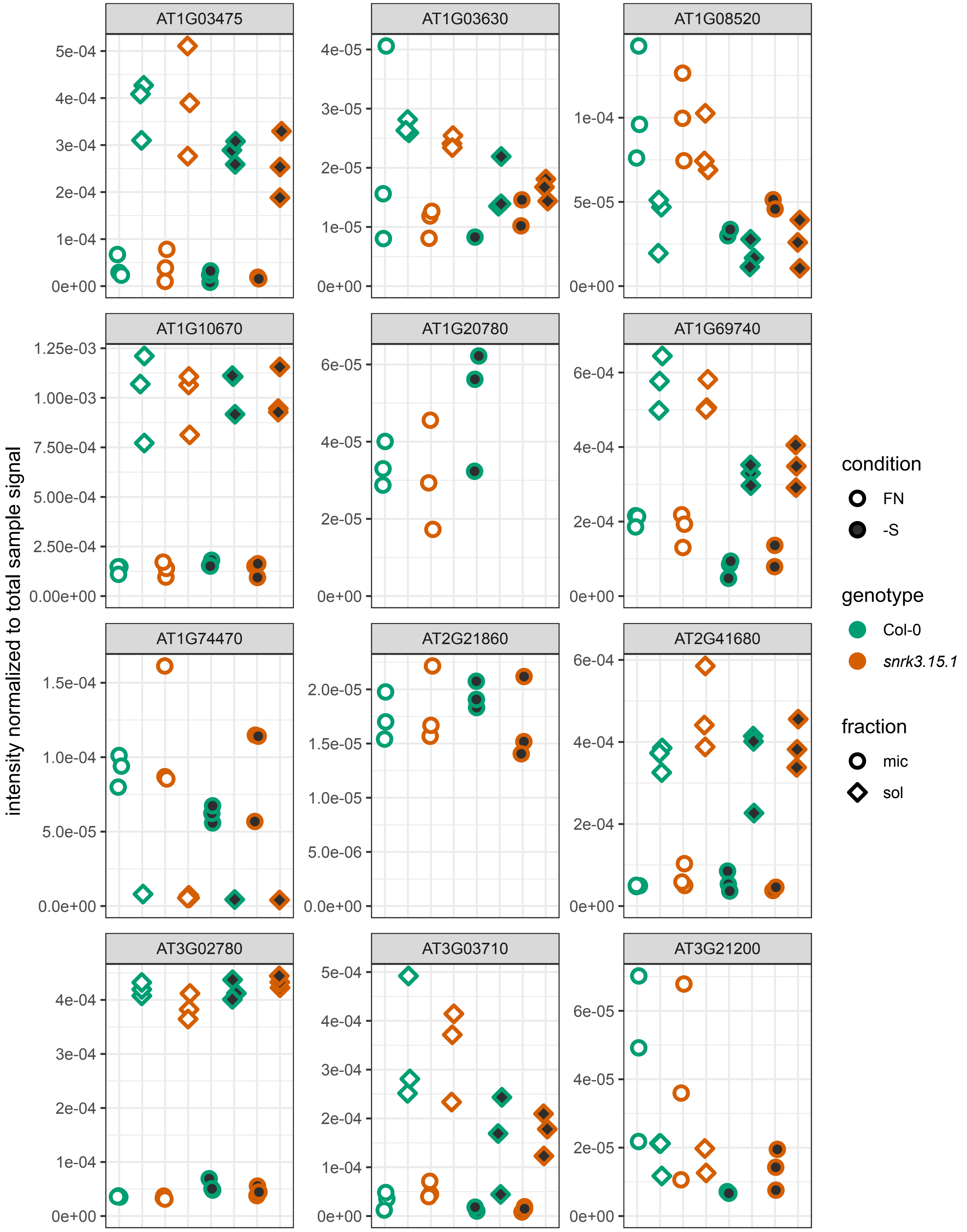

proteins annotated to GO terms positively associated with chlorophyll levels
page 2 of 3

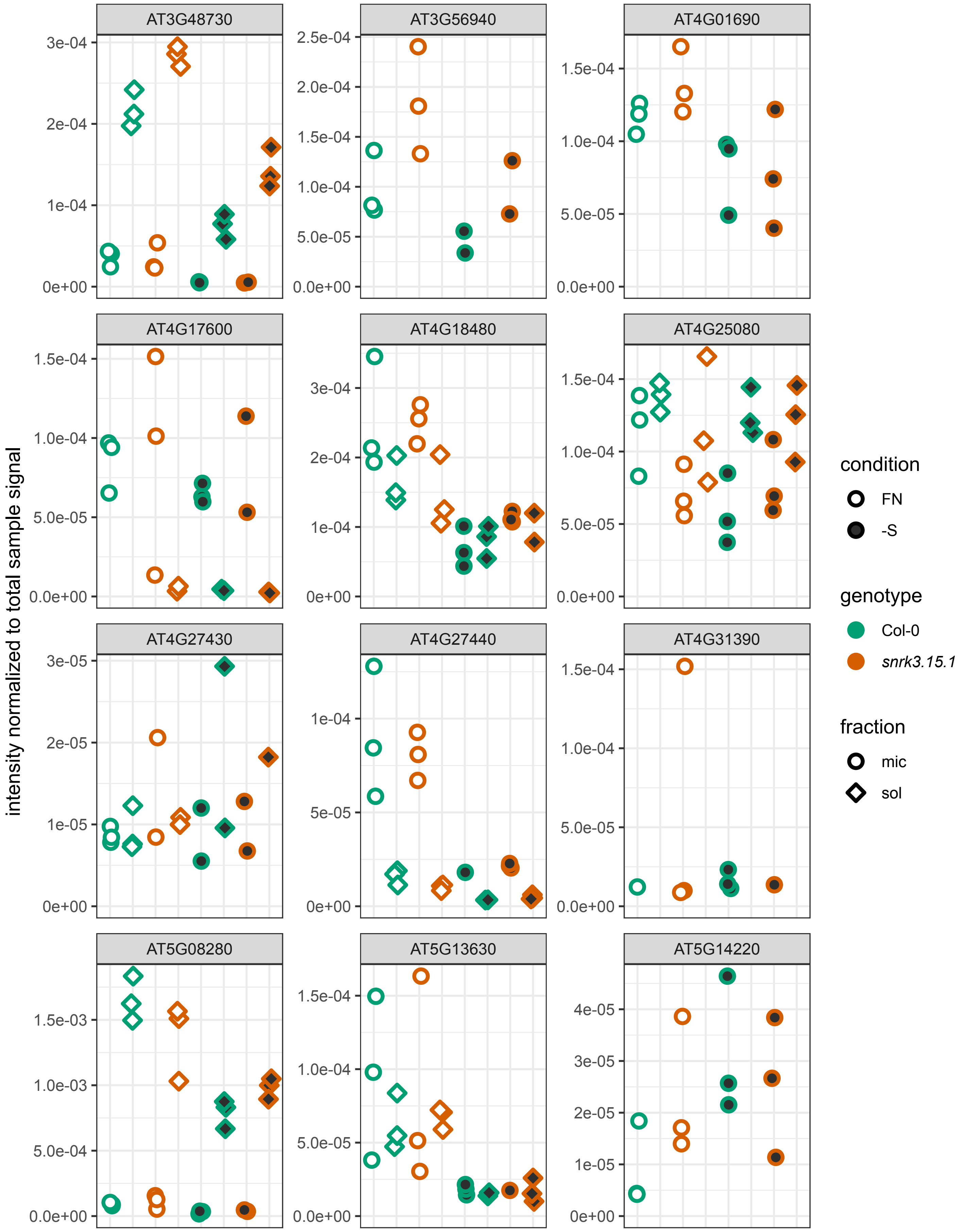

proteins annotated to GO terms positively associated with chlorophyll levels
page 3 of 3

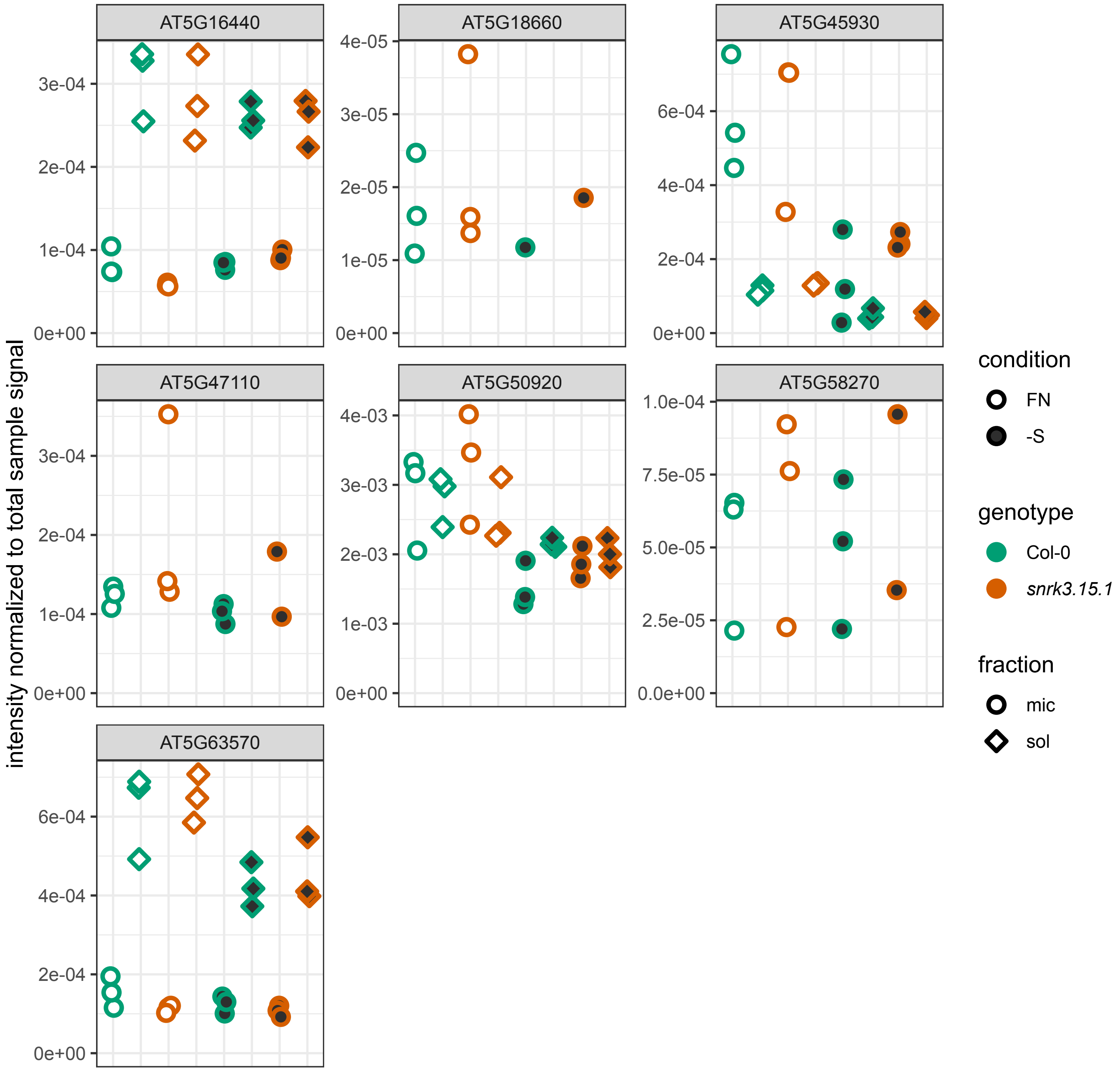

**Supplemental Figure 5**

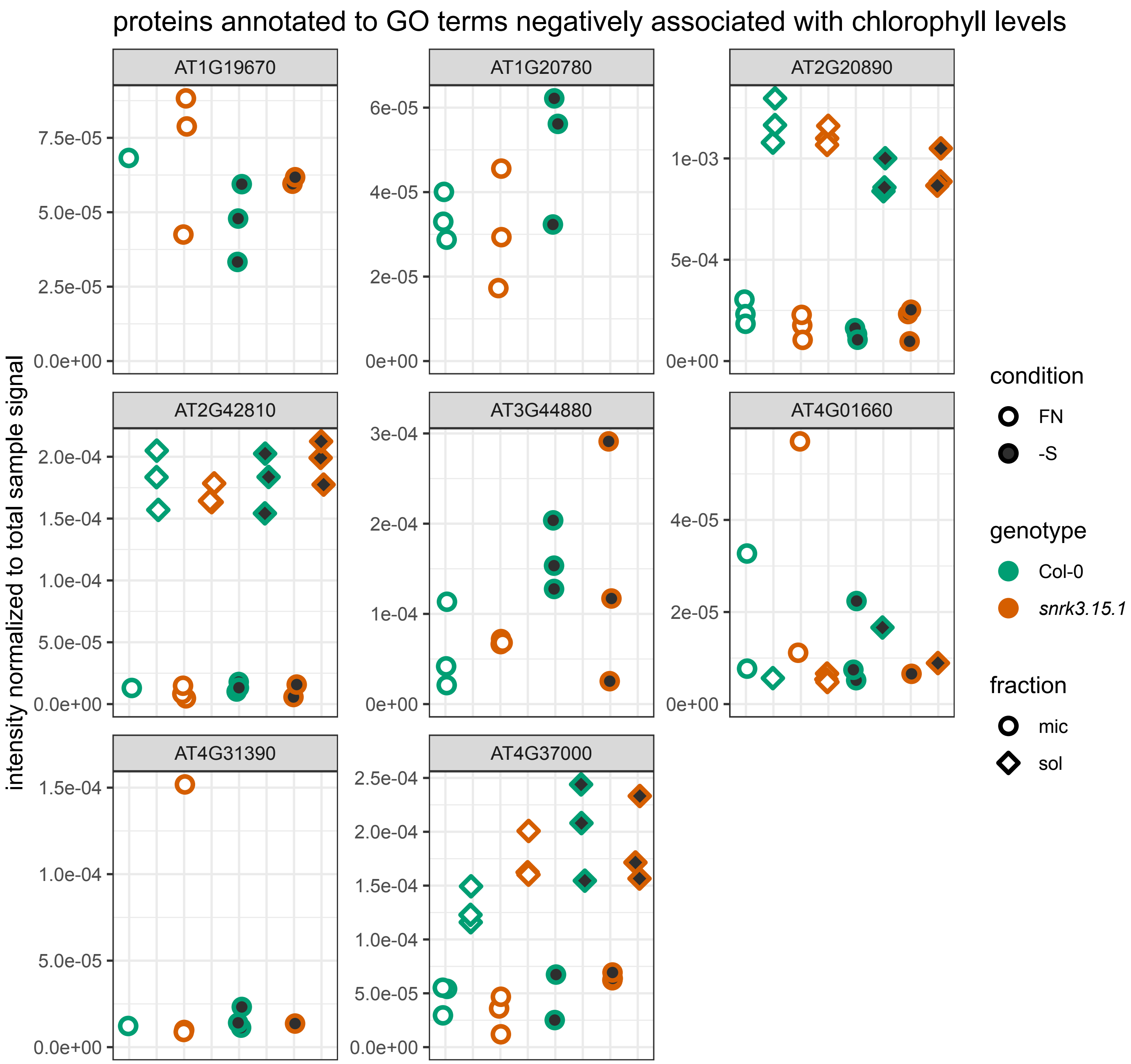

### Supplemental Figure 6

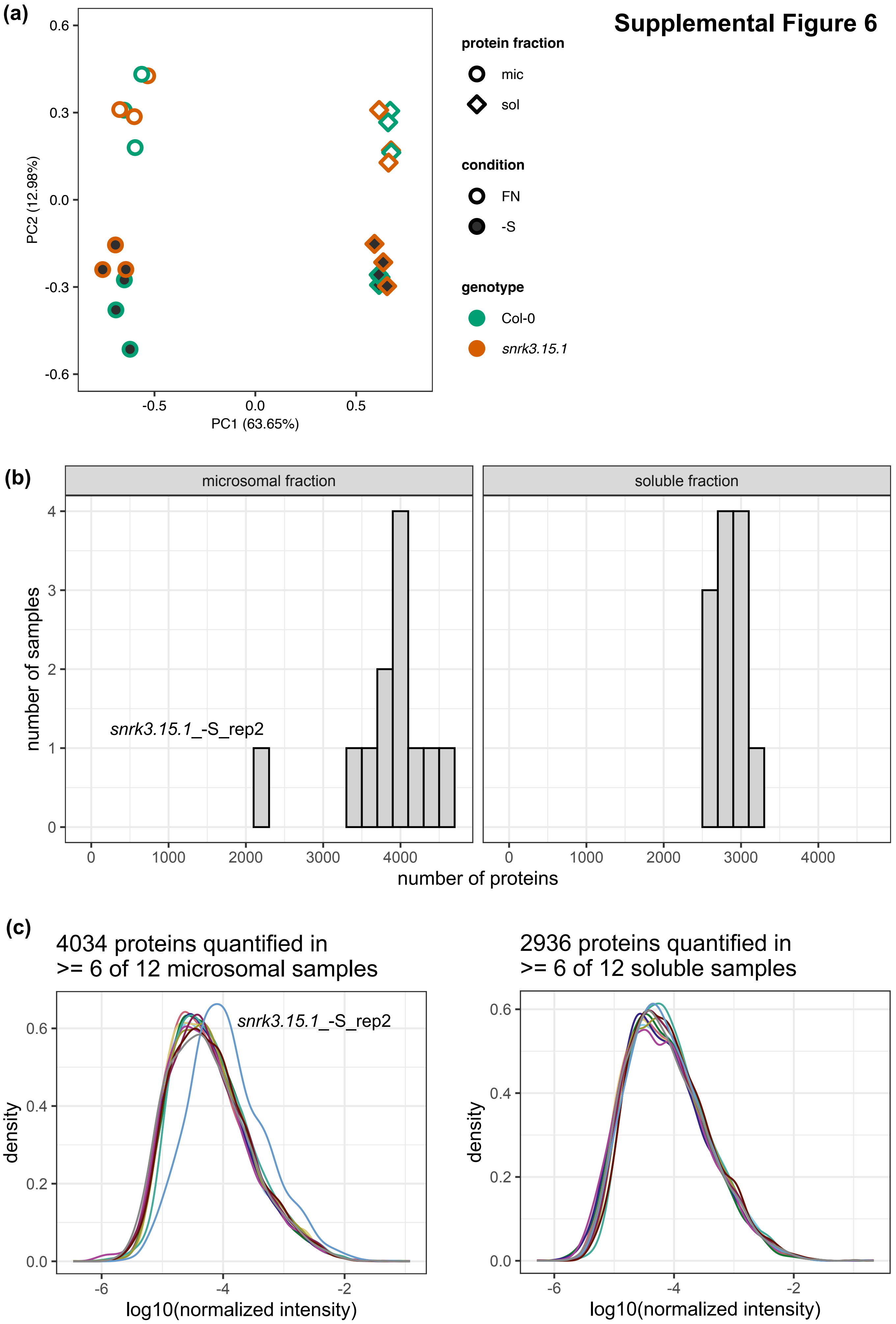

**Supplemental Figure 7**

condition-dependent *snrk3.15*-specific DAPs
(page 1 of 2)

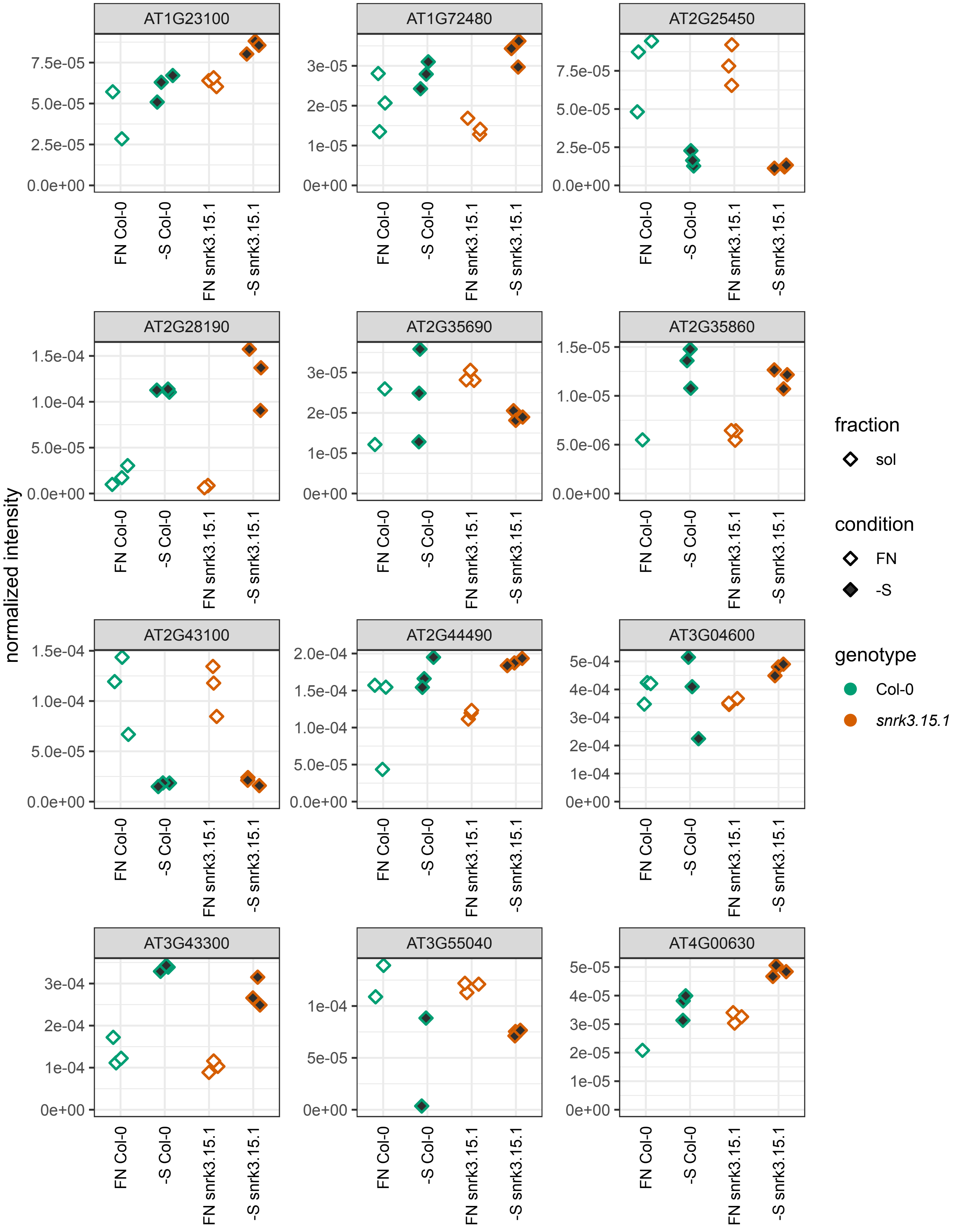

condition-dependent *snrk3.15*-specific DAPs
(page 2 of 2)

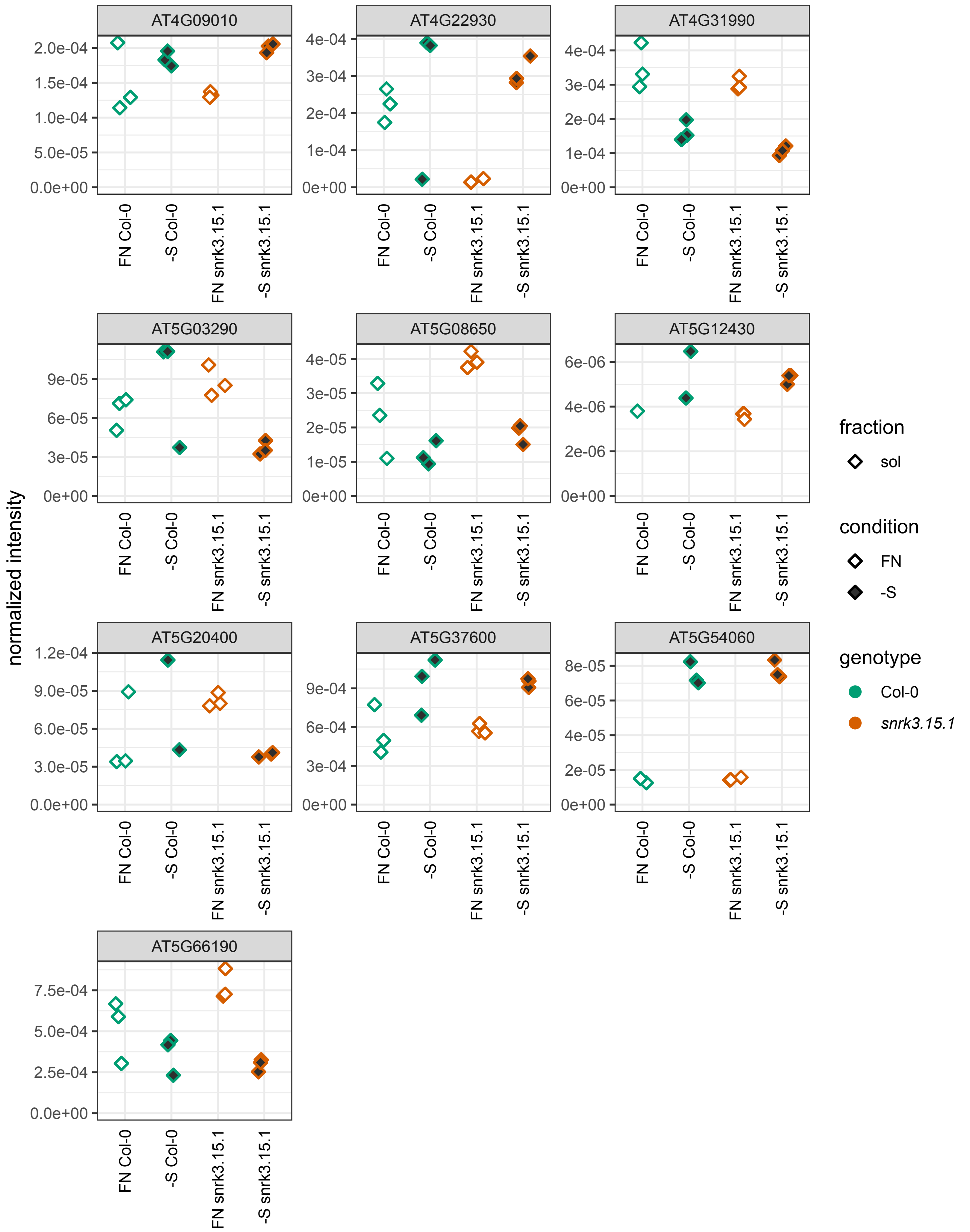

### Supplemental Figure 8

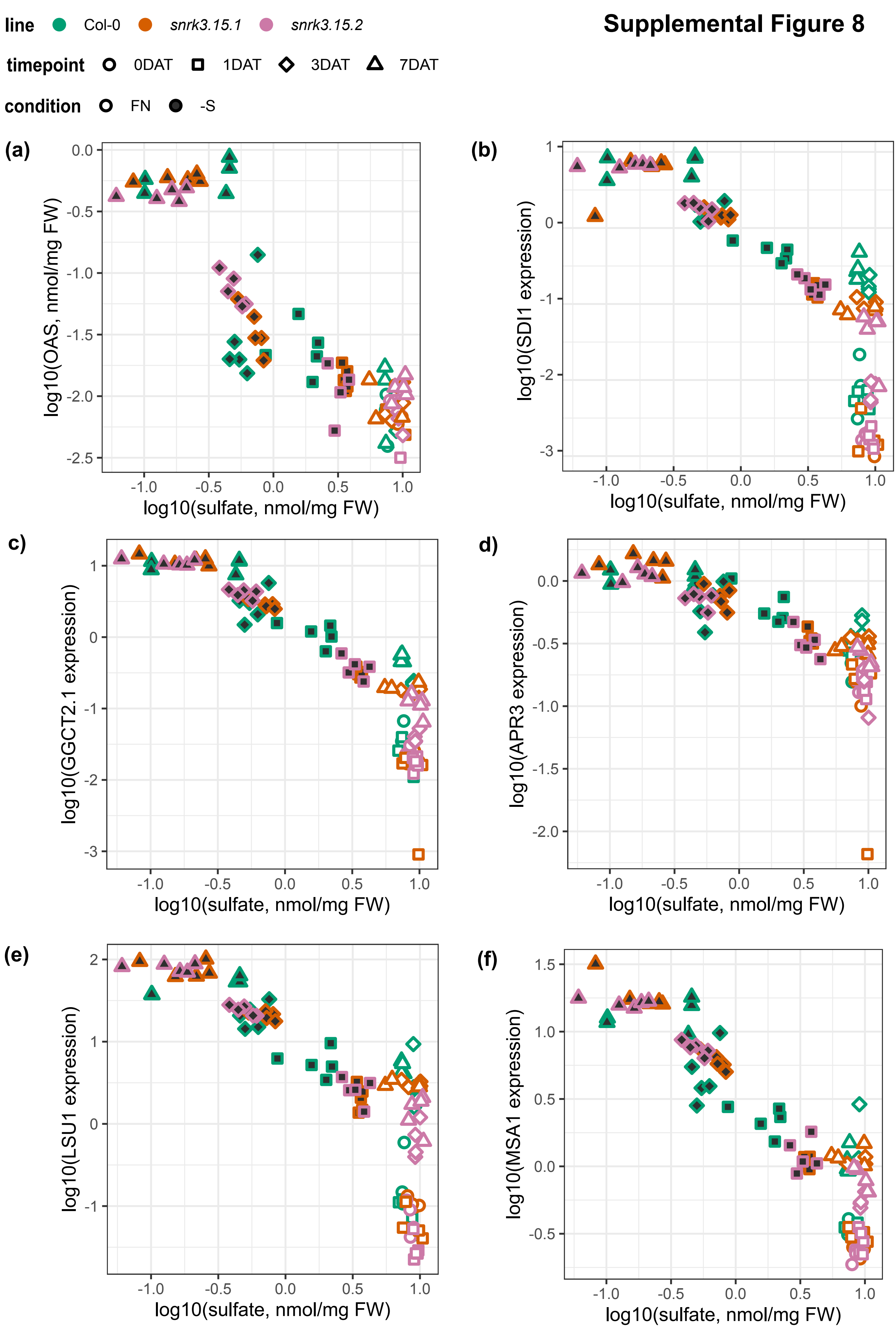
