## Supplemental Table 2 for "*SNRK3.15* is a crucial component of the sulfur deprivation response in *Arabidopsis thaliana*"

### Supplemental Table 2: Media Composition

Composition of modified MS medium. Macro- and micro-elements were added. MS agarose plates were made with either full sulfur supply (+S; 0.75 mM $MgSO_4$) or no sulfur supply (-S; 0.75 mM $MgCl_2$) using a low sulfur (≤ 0.15% sulfate) agarose (Biozym LE Agarose, Biozym Scientific GmbH, Hessisch Oldendorf, Germany).

| **Macroelements** | **Final concentration** |
|---|---|
| $Ca(NO_3)2 \bullet 4H_2O$ | 1.5 mM |
| $KNO_3$ | 1 mM |
| $KH_2PO_4$ | 0.75 mM |
| $MgSO_4 \bullet 7H_2O$ (for FN) | 0.75 mM |
| Fe-EDTA | 1 mM |
| $MgCl_2 \bullet 6H_2O$ (for –S) | 0.75 mM |
| **Microelements** | |
| $MnCl_2 \bullet 4H_2O$ | 10 µM |
| $H_3BO_3$ | 50 µM |
| $ZnCl_2$ | 1.75 µM |
| $CuCl_2$ | 0.5 µM |
| $Na_2MoO_4$ | 0.8 µM |
| KI | 1 µM |
| $CoCl_2 \bullet 6H_2O$ | 0.1 µM |
| **Additives** | |
| MES hydrate | 0.8 g/L |
| Low EEO agarose | 8 g/L |
| Sucrose | 10 g / L |
| pH 5.7 with KOH | |
