## Supplemental Table 3 for "*SNRK3.15* is a crucial component of the sulfur deprivation response in *Arabidopsis thaliana*"

### Supplemental Table 3: qPCR parameters

Sample preparation for qRT-PCR. Reagent volume is written on the table below.

| **Reagent** | **Volume in RT-PCR reaction (5 µl)** |
|---|---|
| cDNA sample (1:9) | 0.5 µl |
| SYBR Green PCR Master Mix | 2.5 µl |
| Primer mix (F + R) 1µM | 2 µl |

PCR program for qPCR.

| | **qPCR cycle** | |
|---|---|---|
| | 50 $^{o}$C 2 min | |
| Initialization | 95 $^{o}$C 10 min | |
| Denaturation<br>Primer annealing and extension | 95 $^{o}$C 15 sec<br>60 $^{o}$C 30 sec | 40 times |
| | 95 $^{o}$C 15 sec | |
| Dissociation | 60 $^{o}$C 15 sec -> 95 $^{o}$C 15 sec | |
